# Ca^2+^ signature gates early auxin signaling in Arabidopsis roots

**DOI:** 10.1101/2025.10.04.680446

**Authors:** Huifang Song, Angel Baudon, Matthias Freund, Marek Randuch, Aleš Pěnčík, Novák Ondřej, Zhaohui He, Kerstin Kaufmann, Nan Li, Matthew Gilliham, Jiří Friml, Rainer Hedrich, Shouguang Huang

## Abstract

Plants must continually balance growth with arrest, especially under stress. Auxin signaling acts as a central regulatory hub in this process, yet the mechanisms that dynamically tune early auxin signalling in real time remain unknown. Here, we used the light-gated, Ca^2+^-permeable Channelrhodopsin 2 variant XXM2.0 to optogenetically impose defined Ca^2+^ signatures on Arabidopsis root cells. Repetitive light activation triggered cytosolic Ca^2+^ signals that in turn suppressed auxin-induced membrane depolarization and Ca^2+^ transients. Moreover, prolonged optogenetic Ca^2+^ stimulation affects auxin-responsive transcriptional reprogramming. As phenotypic output, reversible inhibition of root growth by suppressing cell division and elongation was observed. We further identify a candidate CaM7–CNGC14 module that likely mediates Ca^2+^-dependent gating of early auxin signalling. Our study thus introduces a synthetic biology approach to decompose calcium–auxin crosstalk in plant cells, and demonstrates that optogenetically imposed cytosolic Ca^2+^ signals act as dynamic regulators of auxin susceptibility in roots.

---

The root anchors the plant in the soil and is vital for water and nutrient acquisition. Growth of this underground organ is tightly regulated by the phytohormone auxin, which promotes cell division and elongation via plasma membrane-associated processes as well as transcriptional and translational signaling cascades (*1–3*). In their natural environment, roots frequently encounter stressors such as nutrient fluctuations, dehydration, mechanical damage or pathogen attack. During stress episodes, plants need to decide when to grow and when to pause and instead follow survival strategies.

It has been suggested that plants fine-tune their growth-survival decisions by transiently gating hormone sensitivity (*4–6*). Among candidate regulators, calcium ions (Ca^2+^) are especially compelling, as universal second messengers, associated with early stress signaling in response to cold, salt/osmotic shock, and wounding (*7–12*). In roots, auxin itself can rapidly trigger cytosolic Ca^2+^ elevations, plasma membrane depolarization, and apoplastic alkalinization, collectively contributing to rapid inhibition of root elongation (*1, 2, 13–18*). These fast responses occur within seconds to minutes and depend on the auxin influx carrier AUX1, the predominantly cytoplasmic auxin receptor AFB1, and the Ca^2+^-permeable channel CNGC14 (*13–16, 19*). Intriguingly, auxin-induced Ca^2+^ transients in root hairs are also impaired in *tir1/afb2/afb3* mutant, implicating canonical auxin signalling components in rapid Ca^2+^ responses despite their largely nuclear localization (*13*). How these distinct auxin signalling modules converge to control Ca^2+^ signalling remains unresolved.

A Ca^2+^-dependent feedback circuit within the AUX1–TIR1/AFB–CNGC14 pathway has previously been proposed, based on observations that AUX1-mediated auxin transport is impaired by Ca^2+^ channel inhibition and in *cngc14* mutant, suggesting that elevated cytosolic Ca^2+^ may negatively regulate auxin transport or Ca^2+^ entry itself (*13*). If auxin-induced Ca^2+^ elevations can feed back to attenuate auxin signalling, a similar regulatory logic may also operate during environmental stress, where cold, wounding, or osmotic stimuli generate rapid cytosolic Ca^2+^ transients. In this scenario, stress-induced Ca^2+^ signals could act as a shared second messenger to dynamically tune auxin signaling and transiently prioritize survival over growth. However, direct evidence that cytosolic Ca^2+^ can gate auxin signaling in intact roots remains untested. Addressing these questions has been challenging because, so far, plant Ca^2+^ signals have been mostly provoked by chemical probes (*20, 21*) and/or by modulation of the plasma membrane potential, for example through rapid changes in the composition of the extracellular solution (*22, 23*). However, these approaches are often invasive, prone to off-target effects, or inevitably affect the plant cell plasma membrane potential. This directly impacts ion transport and membrane biology, thereby hindering the ability to address the aforementioned questions.

To overcome these limitations, we in this study used an optogenetics approach to impose well-defined cytosolic Ca^2+^ signals in living root cells. Channelrhodopsins (ChRs) are light-gated ion channels originally derived from algae, and have been adapted as powerful tools for manipulating membrane potential and ion fluxes in plant cells (*24–29*). Here, we expressed the highly Ca^2+^-permeable ChR2 variant XXM2.0 (*26, 27*) in Arabidopsis roots to generate spatially and temporally controlled Ca^2+^ signatures. This approach enabled us to study how Ca^2+^ modulates auxin biology over both short and long timescales. By combining optogenetics, electrophysiology, live-cell imaging, transcriptomics, and developmental phenotyping, we show that Ca^2+^ functions as a dynamic regulator of early auxin signaling.

## ChRs enable non-invasive control of electrical and Ca^2+^ signals in root cells

In Arabidopsis roots, auxin application rapidly triggers membrane depolarization and cytosolic Ca^2+^ transients, both of which are thought to contribute to downstream signaling and developmental responses (*13, 18*). Yet, how these two signals interact and how each contributes to auxin output remains unclear. Light-gated ion channels provide a non-invasive approach to study these dynamics in living tissues (*30, 31*), particularly in roots, which act as multicellular sensory organs integrating environmental signals via Ca^2+^ and electrical signals (*32, 33*).. We expressed the light-gated anion channel GtACR1 and Ca^2+^-permeable ChannelRhodopsin 2 variant XXM2.0 under the UBQ10 promoter in Arabidopsis. All constructs were C-terminally fused to eYFP for subcellular localization analysis and associated with a retinal-generating dioxygenase (see Methods for technical details). Confocal microscopy showed that both GtACR1 and XXM2.0 localized predominantly to the plasma membrane of root cells, confirming correct and stable expression (Fig. 1A).

**Fig. 1.**
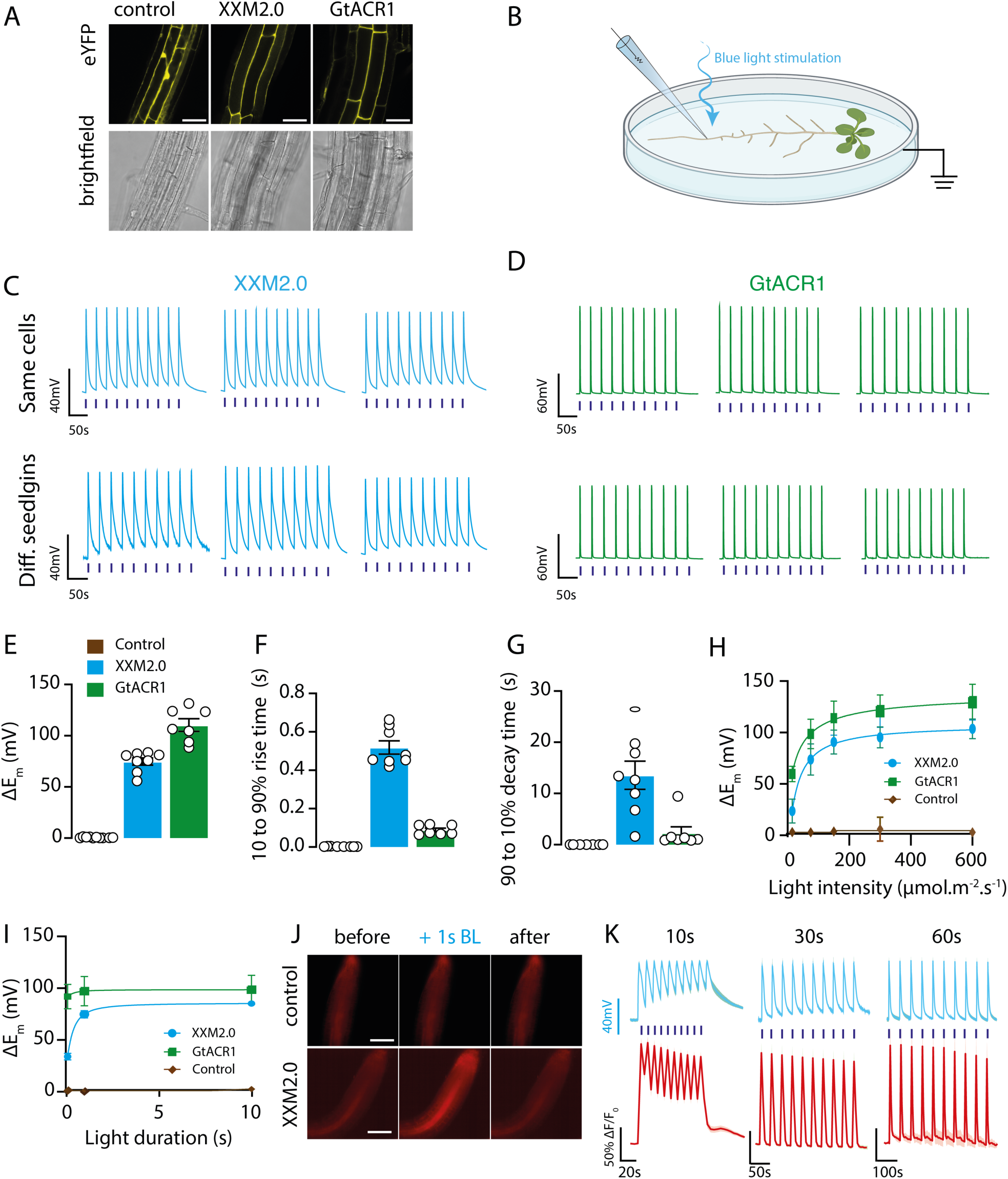
High-fidelity electrical signals induced by XXM2.0 and GtACR1 in the root. (**A**) Confocal images of root cells expressing control, XXM2.0, or GtACR1. Scale bars represent 50 μm. (**B**) Schematic illustration of light stimulation and electrophysiological recording in Arabidopsis seedlings. (**C-D**) Membrane potential changes evoked by trains of BL pulses (1 s pulse, 30 s interval, 10 total stimuli) in seedlings expressing XXM2.0 (C) or GtACR1 (D). Upper traces represent recordings from the same cells of a seedling; lower traces show responses from three different seedlings. (**E-G**) Quantification of depolarization amplitude, rise time, and decay time in response to photostimulation in seedlings expressing XXM2.0 (blue, n = 8), GtACR1 (green, n = 7), or the control (brown, n = 11). (**H–I**) Quantification of the effects of light intensity (H) and light duration (I) on membrane depolarization. Blue = XXM2.0 (n = 8), Green = GtACR1 (n = 8–9), Brown = control (n = 11–13). All data are shown as mean ± SEM. (**J**) Representative images of R-GECO1 signals in response to 1-s BL stimulation in the control and XXM2.0-expressing seedlings. (**K**) Averaged depolarization (blue) and Ca^2+^ signal (red) traces, evoked by light stimulation with varying inter-pulse intervals in XXM2.0 expressing seedling.

To evaluate whether XXM2.0 and GtACR1 reliably trigger electrical signals in root cells, we performed intracellular recordings using sharp microelectrodes in the meristematic zone of 3-5-day-old Arabidopsis seedlings (Fig. 1B). Upon blue light stimulation (1 s pulse, 100 µE), root cells expressing either XXM2.0 or GtACR1 became electrically excited (Fig. 1C–D). When seedlings were exposed to a series of repetitive light pulses, both light-gated channels triggered robust, repetitive membrane depolarizations, and this effect was reproducible within the same cell and between different seedlings (Fig. 1C–D). This demonstrates that using GtACR1 and XXM2.0-based optogenetic tools, membrane potential changes can be precisely and uniformly controlled at the cell, tissue, and population level.

Under 470 nm light at 150 µE intensity, a single 1-s pulse induced rapid depolarization in root cells: ∼110 mV for GtACR1 and ∼75 mV for XXM2.0. In contrast, no significant change was observed in the control seedlings (Fig. 1E-G). The magnitude of depolarization was strongly dependent on both light intensity and pulse duration (Fig. 1H-I). Interestingly, the two optogenetic tools displayed unique temporal dynamics. In GtACR1-expressing cells, the membrane potential precisely followed the light pulse with rapid rise (tau= 89.15 ± 0.95 ms) and decay (tau= 2.29 ± 1.20 s) phases, mirroring the BL stimulation kinetics (Fig. 1D and Fig. 1F-G). By contrast, XXM2.0-induced depolarizations exhibited a much slower recovery (tau= 13.56 ± 2.75 s) (Fig. 1C, Fig. 1F-G). This delay in the XXM2.0 -evoked electrical response relative to the Ca^2+^ transient likely reflects the time required to biochemically activate downstream depolarizing channels - such as endogenous S- and R-type anion channels (*20, 26*) - upon Ca^2+^ elevation, and to deactivate them upon stimulus cessation.

To investigate how the temporal pattern of XXM2.0 stimulation affects root electrical and Ca^2+^ responses, we co-expressed the XXM2.0 with the genetically encoded Ca^2+^ indicator R-GECO1 and applied repetitive 1-s BL pulses at varying intervals (10 s, 30 s, and 60 s). At 30- and 60-s intervals, both Ca^2+^ transients and membrane depolarizations were well separated, returning to baseline between consecutive light pulses (Fig. 1J-K). However, stimulating the plants every 10 s led to a temporal summation of the signals, indicating that cells need more time to return to the resting state (Fig. 1K).

## Optogenetic Ca^2+^ priming suppresses auxin-induced membrane excitation

GtACR1 and XXM2.0 enable precise control of membrane voltage and Ca^2+^ in root cells. We next used these tools to test whether prior electrical or Ca^2+^ stimulation modulates auxin response. Meristematic root cortical cells were impaled with voltage-recording microelectrodes, and IAA was locally applied through a glass micropipette as a brief 5-s pulse (“IAA puff”) (Fig. 2A). This is because auxin-induced membrane depolarization is tightly coupled to rapid auxin perception and displays dose dependence (*13, 19, 34*), we used it here as a quantitative readout of the fast auxin response. Control roots depolarized in response to IAA puffs but not to mock (Fig. 2B-C). Following the IAA pulse, the membrane potential progressively depolarized, reaching its 40-60 mV peak within 60 s, then returned toward the pre-stimulation baseline over the next 300 s (Fig. 2C-E). Although the first IAA-induced depolarization was similar between control and XXM2.0-expressing seedlings, the repetitive blue light stimulation substantially reduced the depolarization amplitude triggered by the second IAA application in XXM2.0 (60% in Fig. 2C-E) but not in control seedlings. In contrast, the second IAA response remained unchanged in both amplitude and kinetics in GtACR1-expressing roots (Fig. 2C-E). This finding indicates that Ca^2+^ influx, rather than membrane depolarization alone, is responsible for attenuating the membrane auxin response. The IAA-induced membrane depolarization response recovered when more time was allowed between termination of XXM2.0-induced Ca^2+^ signals and the second IAA application (Fig. S1). To define recovery kinetics, we monitored IAA-evoked depolarization at 30–300 s after a 20-pulse light train and found that was fully restored within ∼100 s (Fig. 2F). To test whether Ca^2+^-dependent modulation of the rapid auxin response scales with the stimuli applied, we varied the number of BL pulses applied to XXM2.0 roots. As few as three BL pulses already reduced auxin-induced membrane depolarization to 80% of the initial response, with progressively stronger inhibition with 10 pulses (Fig. 2G-I). Auxin responses fell below 50% of baseline after 20 pulses (Fig. 2I), suggesting that the imposed Ca^2+^ dose determines the degree to which auxin-induced membrane depolarization is suppressed. This notion was supported by direct Ca^2+^ measurements. R-GECO1-based Ca^2+^ imaging in XXM2.0 × R-GECO1 seedlings showed that auxin-induced Ca^2+^ transients were strongly suppressed after BL stimulation (Fig. 2J-L), a result well in line with the root membrane electrical response.

**Fig. 2.**
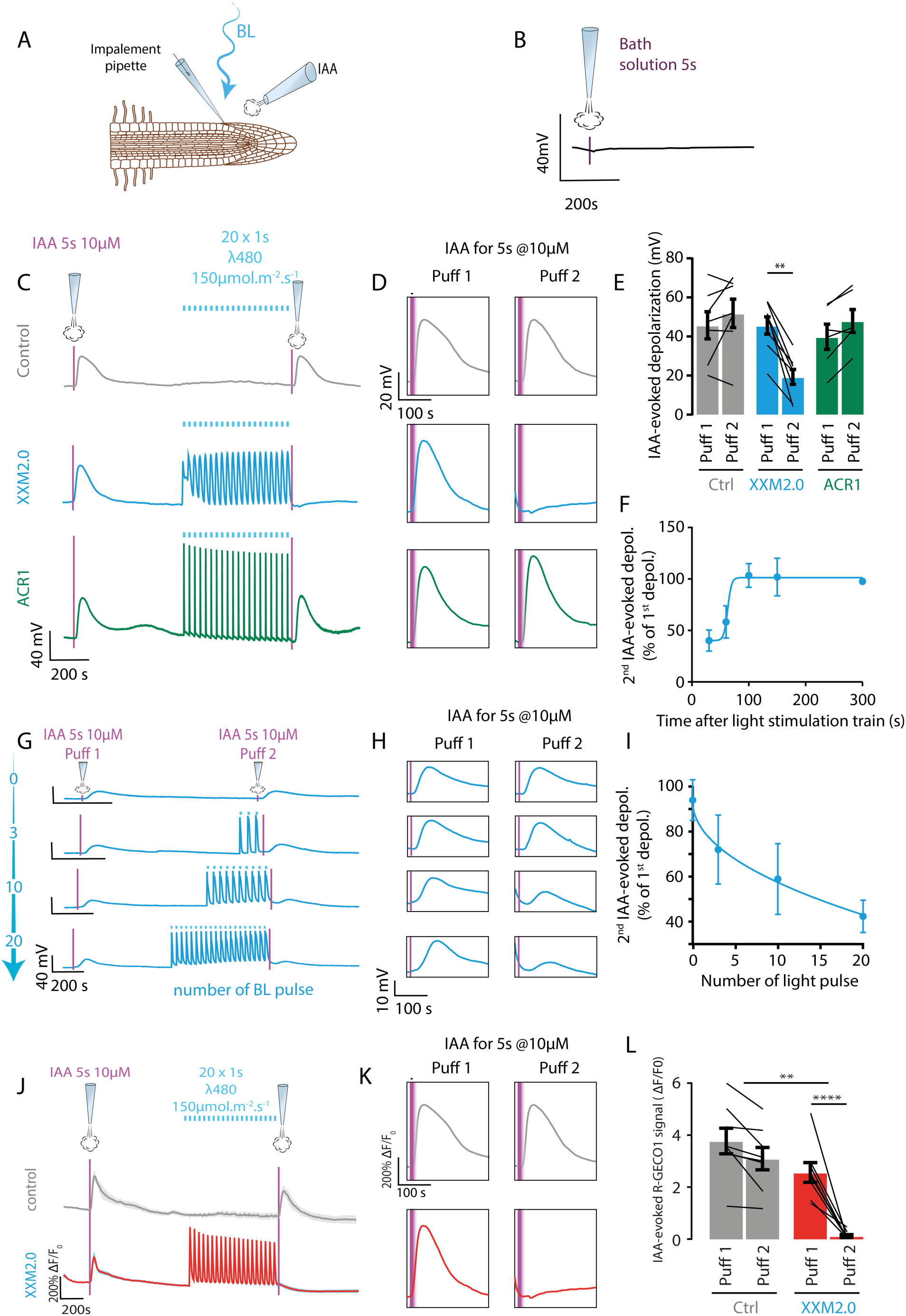
XXM2.0 photostimulation alters early auxin signalling in roots. (**A**) Schematic illustration of the combined auxin application and electrophysiological recording setup. IAA (10 µM in the pipette tip) was locally applied for 5-s using a glass micropipette (diameter = 10 µm). (**B**) Membrane potential trace of a root before and after the local application of bath solution (marked with the purple line). (**C**) Membrane potential of cortical cells in the root meristematic zone during a first local application of IAA (Puff 1), a train of light stimulation (ʎ = 470nm, 150 µE, 20 pulses of 1 second every 30 seconds), and a second local IAA application (Puff 2) in seedlings expressing the control (grey), XXM2.0 (blue), or GtACR1 (green). Purple arrows show IAA application times. (**D**) Zooms on IAA-evoked depolarization shown in C. (**E**) Quantification of IAA-evoked depolarization before and after the train of light pulses in the control (grey, n=7), XXM2.0 (blue, n=8), or GtACR1 (green, n=6)-expressing cells. (**F**) Relative IAA-evoked depolarisation amplitude 30s, 50s, 100s, 150s, and 300s after the last photostimulation of a 20-pulse light train (n=6, 5, 4, 4, and 3, respectively). (**G-H**) Representative membrane recording traces (G) and Insets (H) showing the depolarization triggered by local IAA applications before (left, Puff 1), or after 0, 3, 10, or 20 light pulses (right, Puff 2). Data in each row comes from the same cell. (**I**) Relative IAA-evoked depolarisation amplitude after 0, 3, 10, or 20 light stimulations (n=5, 6, 9, and 8, respectively). (**J**) Ca^2+^ signals in root cells during the first local application of IAA, a train of light stimulation, and a second local IAA application in seedlings expressing either the control (gray) or XXM2.0 (red). (**K**) Zoom-in of the IAA-evoked Ca^2+^ signal shown in J. (**L**) Average of IAA-evoked Ca^2+^ signal before and after the train of light pulses in the control (grey, n=7) and XXM2.0 (blue, n=8)-expressing cells. ** *p* < 0.01, **** *p* < 0.0001.

To ask whether Ca^2+^-dependent attenuation of subsequent auxin responses depends on specific features of the Ca^2+^ signature, we systematically altered the XXM2.0 stimulation protocol before applying a second IAA pulse. We first tested whether a single sustained Ca^2+^ elevation is sufficient. A 20-s blue-light pulse strongly suppressed the second IAA-induced Ca^2+^ response, producing an effect comparable to repetitive stimulation (Fig. S2A). We next varied stimulation interval (10 to 60 s), light intensity (75 to 300 µmol·m^-2^·s^-1^), and pulse duration (0.1 to 2 s) to generate distinct Ca^2+^ dynamics. Across all tested conditions, subsequent auxin responses remained strongly attenuated (Fig. S2B–D). Thus, within the tested parameter range, diverse Ca^2+^ stimulation patterns converged on a similar attenuation of subsequent auxin responses in root meristem cells.

To test whether physiological Ca^2+^ signals similarly modulate subsequent auxin responses, we compared the effects of XXM2.0-evoked Ca^2+^ signals with those provoked after cold and glutamate pre-treatment. Cold treatment is known to reproducibly excite the plasma membrane and trigger cytosolic Ca^2+^ transients (*35–37*). To test if cold-induced Ca^2+^ signals affect the subsequent auxin-induced Ca^2+^ response, IAA was applied immediately after the cold-induced Ca^2+^ signal returned to baseline in root cells. As shown in Fig. 3A-B, cold treatment markedly suppressed the second IAA-induced R-GECO1 Ca^2+^ signal by ∼60% compared to the initial. After a recovery period of 15 – 20 mins, however, a subsequent IAA puff elicited a response reaching ∼90% of the initial amplitude, indicating that the auxin-induced Ca^2+^ response was restored. Similarly, glutamate, a classical wounding-associated chemical signal, induced a strong Ca^2+^ transient that suppressed response to an auxin stimulation by ∼75% (Fig. 3C-D). Like the cold response, the auxin response fully recovered after a recovery period of 15-20 min. Based on the findings above, we assumed that auxin-induced Ca^2+^ signals themselves might create negative feedback on subsequent auxin responses. Indeed, a second IAA pulse delivered 180 s after the first one provoked a markedly weaker response (Fig. 3E-F). However, similar to the cold- and glutamate-induced responses, the auxin-induced Ca^2+^ response recovered to ∼85% following a 10 min recovery period. Consistently, when membrane depolarization was used as an independent readout, a second IAA application delivered after an initial IAA pulse elicited a markedly weaker response, decreasing from ∼40 mV to ∼20 mV (Fig. 3G–H), an attenuation highly comparable to that induced by XXM pre-stimulation (Fig. 2E). Thus, cytoplasmic Ca^2+^ transients generated by the hormone itself or stressors appear to limit these rapid auxin responses.

**Fig. 3.**
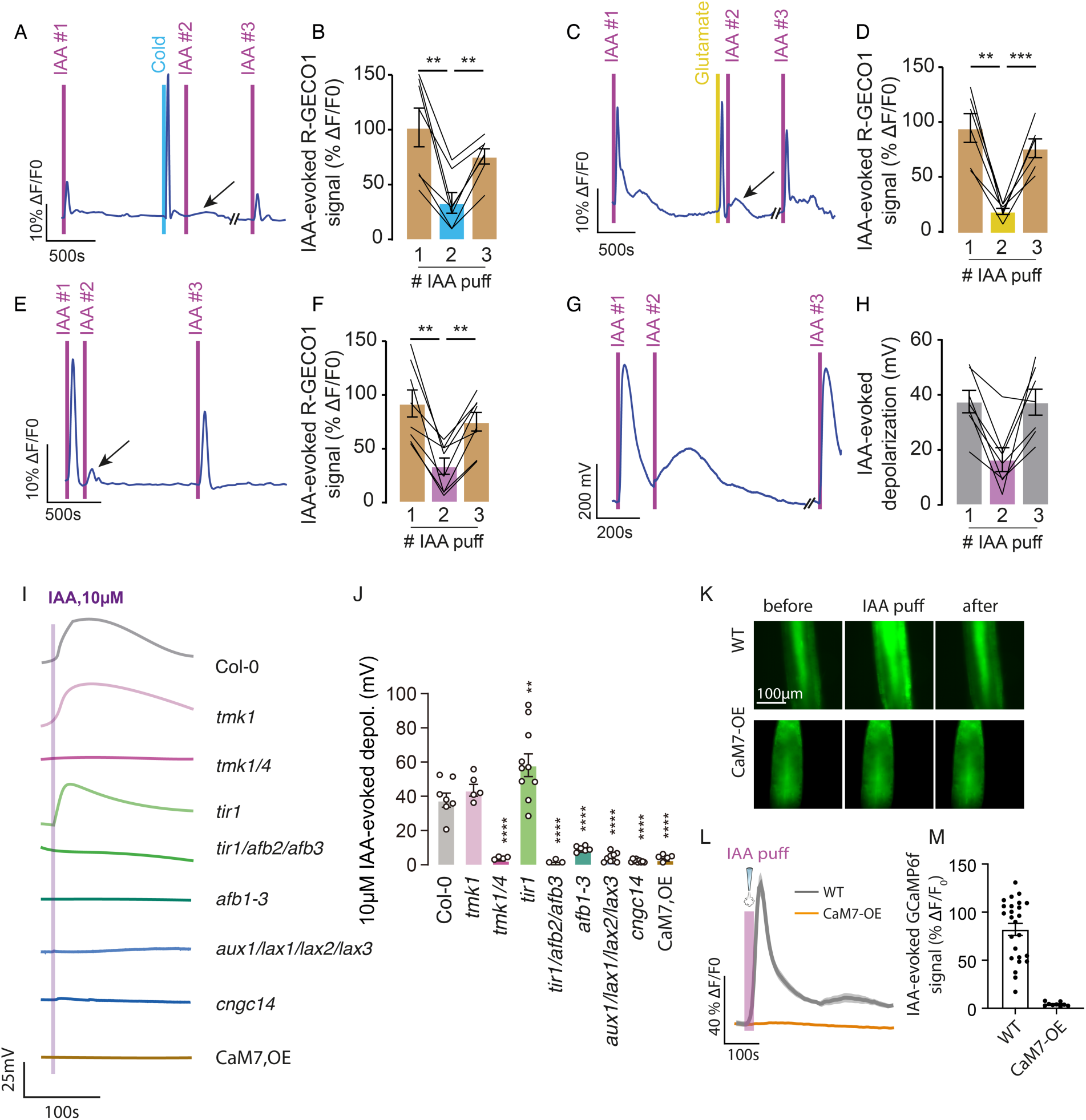
Ca^2+^ transiently suppresses auxin-induced responses and implicates a CaM7– CNGC14 module. (A, C, E) Representative R-GECO1 fluorescence traces showing IAA-evoked cytosolic Ca^2+^ responses before and after cold shock (cyan bar; A), 5 mM glutamate treatment (yellow bar; C), or a following IAA application (magenta bars; E) in Arabidopsis root meristematic cells. Arrows indicate attenuated second IAA responses. (B, D, F) Quantification of IAA-evoked R-GECO1 fluorescence increases (ΔF/F0) following cold treatment (B), glutamate treatment (D), or repeated IAA stimulation (F). Each line represents one biological replicate. Bars indicate mean ± SE (n = 7, 6, and 8, respectively). (G–H) Representative membrane potential trace (G) and quantification (H) of IAA-evoked depolarization following sequential IAA puff in root cortical cells. Each line represents one biological replicate. Bars indicate mean ± SE (n = 9). (I) Representative membrane potential traces evoked by local application of 10 μM IAA in different auxin-related mutants and CaM7-overexpression (CaM7-OE) plants. The purple line indicates the timing of IAA application. (J) Quantification of IAA-evoked membrane depolarization shown in panel (I). Each dot represents one biological replicate. Bars indicate mean ± SE. (K–M) Representative GCaMP6f fluorescence images (K), fluorescence traces (L), and quantification (M) of auxin-induced cytosolic Ca^2+^ responses in wild type (Col-0/GCaMP6f) and CaM7-OE x GCaMP6f roots following IAA puff. Scale bar, 100 μm. Bars indicate mean ± SE; each dot represents one biological replicate. ** *p* < 0.01, *** *p* < 0.001.

## Ca^2+^-regulated early auxin signaling implicates a CaM7–CNGC14 feedback module

To determine whether Ca^2+^-dependent gating of rapid auxin responses relies on key components of the auxin signaling network, we analyzed membrane potential changes in Arabidopsis mutants defective in auxin transport, perception, or Ca^2+^ signaling. In response to 10 µM IAA, wild-type roots displayed a pronounced membrane depolarization of ∼35–40 mV, whereas several canonical auxin signaling mutants, including *aux1/lax1/lax2/lax3, afb1*, and *cngc14*, showed strongly reduced or nearly absent responses (∼0–5 mV, Fig. 3I-J). These results confirm the importance of AUX/LAX-mediated auxin influx, AFB1-dependent perception, and CNGC14-mediated Ca^2+^ signaling for normal auxin responsiveness. By contrast, *tmk1* retained a substantial IAA-induced depolarization (∼40 mV), whereas the *tmk1/4* double mutant, which disrupts a non-canonical auxin signaling branch (*38, 39*), largely lost responsiveness (Fig. 3I-J), indicating differential contributions of canonical and non-canonical auxin signaling branches to the electrical auxin response.

Because CNGC14 was previously suggested to be negatively regulated by an unidentified calmodulin (*13*), we focused on the Ca^2+^ sensor CaM7, which was later shown to inhibit CNGC14 activity in the *Xenopus* oocyte system (*40*). Whether CaM7 contributes to auxin responsiveness in planta, however, remained unknown. We therefore tested the role of CaM7 in auxin signalling. Strikingly, CaM7-overexpression (CaM7-OE) plants phenocopied *cngc14* mutants. Similar to *cngc14*, CaM7-OE roots showed little or no membrane depolarization in response to 10 µM IAA (Fig. 3I–J). Furthermore, IAA-induced cytosolic Ca^2+^ signals were abolished in CaM7-OE plants (Fig. 3K–M). At the developmental level, CaM7-OE roots displayed elongated primary roots resembling the *cngc14* phenotype (Fig. S3). Together, these findings implicate a CaM7–CNGC14 module as a candidate mechanism for Ca^2+^-dependent attenuation of early auxin responses, in which a preceding Ca^2+^ transient may engage CaM7 to suppress CNGC14-mediated signalling.

## Optogenetically-imposed Ca^2+^-transients affect transcriptional auxin signaling in the root tip

Given that Ca^2+^ signals change the auxin response of root cells, we asked if Ca^2+^ affects the overall auxin content of the Arabidopsis seedling roots. Auxin profiling of whole root samples by high-performance liquid chromatography combined with mass spectrometry (LC-MS), however, revealed no significant changes in IAA biosynthesis and/or metabolism following XXM2.0-induced Ca^2+^ stimulation (Fig. S4). Similarly, we found no evidence for XXM2.0-induced Ca^2+^ effects on polar localization of PIN1 and PIN2 auxin transporters in roots (*41*), suggesting that root auxin fluxes remain unaffected (Fig. S5). To assess whether optogenetically imposed Ca^2+^ signals modulate auxin-responsive transcription, we used the *DR5v2*::n3GFP reporter, which sensitively reflects auxin-dependent transcriptional output (*42*). To this end, we crossed the *DR5v2*::n3GFP with XXM2.0 or GtACR1 and monitored fluorescence in root tips using confocal microscopy. Seedlings were grown under RL for 3 days and then subjected to BL (1 s 100 µE pulses every 30 s) for 1-48 h. Under RL conditions, *DR5v2*::n3GFP exhibited strong expression in the columella, quiescent center, meristem, and elongation zone across all genotypes (Fig. 4A-B). In contrast, XXM2.0 roots exposed to BL exhibited a progressive and significant reduction in *DR5* response compared to RL-treated controls (Fig. 4A and C). Quantification revealed that *DR5* intensity in the meristematic and elongation zones of XXM2.0 roots decreased by ∼37 to 50% throughout the 1 to 48 h period (Fig. 4C). This decline was statistically significant at all time points, and was not observed in the controls, where *DR5* response remained stable under the same BL treatment (Fig. 4C). Importantly, in contrast to XXM2.0, GtACR1 roots did not show a reduction in *DR5* signal (Fig. 4A-C). This indicates that suppression of DR5 activity is caused by Ca^2+^ influx, rather than membrane depolarization alone. Notably, switching XXM2.0 seedlings back to RL restored DR5 signal intensity to baseline levels (Fig. 4D-E).

**Fig. 4.**
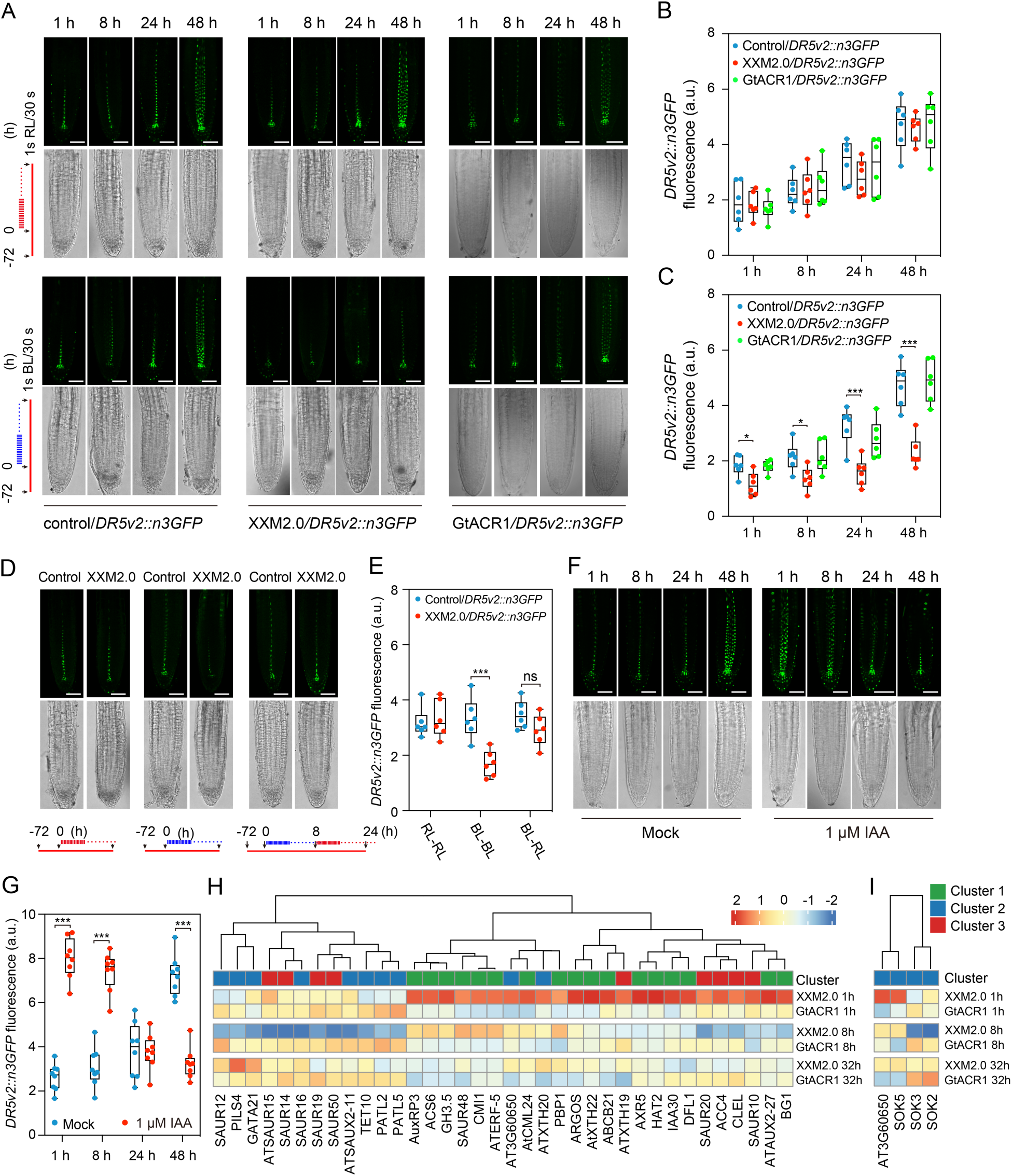
Imposed Ca^2+^ signals are accomapanied by reduced DR5 signaling and reprogramming of auxin-responsive transcripts. (**A**) Time course of *DR5v2::n3GFP* expression in root tips of Col-0/ *DR5v2::n3GFP* (WT) and XXM2.0/*DR5v2::n3GFP* and GtACR1/*DR5v2::n3GFP* seedlings under red light (top) or blue light (bottom) pulse stimulation (1 s every 30 s, 100 µE). Roots were imaged at 1 h, 8 h, 24 h, and 48 h after the onset of stimulation. Maximum intensity projections (green) and corresponding brightfield images are shown. Scale bars = 50 µm. (**B-C**) Quantification of *DR5v2::n3GFP* fluorescence in root tips across time points shown in (A). Under red light (B), reporter signal increases progressively in both genotypes. Under blue light (C), signal accumulation in XXM2.0 seedlings is significantly suppressed compared to Col-0 at 8 h, 24 h, and 48 h. Data represent n = 5-6 roots per group and are presented in arbitrary units (a.u.). (**D-E**) Fluorescence of *DR5v2::n3GFP* in root tips of WT and XXM2.0/*DR5v2::n3GFP* seedlings under pulsed light: 24 h RL (left), 24 h BL (middle), and 8 h BL + 16 h RL (right) (D), and corresponding quantification (E). Data represent n = 5-6 roots per group and are presented in arbitrary units (a.u.). (**F**) Time course imaging of root tips from transgenic Arabidopsis seedlings (*DR5v2*::n3GFP) treated with 1 μM IAA for 1, 8, 24, and 48 h. Seedlings sprayed with the corresponding mock solution were used as controls. Maximum intensity projections (green) and corresponding brightfield images are shown. Scale bars, 50 μm. (**G**) Quantification of *DR5v2*::n3GFP fluorescence intensity in root tips at the indicated time points is shown in (F). Fluorescence was quantified in ImageJ as the mean gray value and was presented in arbitrary units (a.u.). (**H-I**) Heatmap showing the expression patterns of auxin-responsive (F) and root morphogenesis (G) genes across XXM2.0 and GtACR1 roots at 1 h, 8 h, and 32 h post-stimulation. Genes were grouped by hierarchical clustering into three expression clusters (1, 4, and 5; color-coded). Data are mean ± SD. * *p* < 0.05, *** *p* < 0.001, ns = not significant.

To assess whether attenuation of auxin transcriptional output also occurs within the endogenous auxin signaling system, we monitored DR5 responses during prolonged exogenous IAA treatment. Similar to XXM2.0 activation, DR5 reporter intensity progressively declined and became markedly reduced by 24–48 h (Fig. 4F-G). Thus, sustained auxin exposure itself attenuates auxin transcriptional output over time, paralleling the effects of XXM-induced Ca^2+^ stimulation. These findings indicate that cytosolic Ca^2+^ elevation via XXM2.0 does not suppress auxin homeostasis and transport but suppresses the transcriptional auxin output in the root tip, consistent with a role for Ca^2+^ in regulating canonical auxin signaling, although growth arrests associated with altered developmental states cannot be excluded.

## Optogenetic Ca^2+^ stimulation reprograms root transcript landscape

Having shown that activation of XXM2.0 attenuates transcriptional auxin reporter activity, we next asked whether the Ca^2+^ signal reprograms auxin-regulated transcription globally. Although previous studies have shown that auxin-induced Ca^2+^ signals contribute to downstream responses (*13, 18*), the specific role of Ca^2+^ in regulating transcription remains unclear, as auxin simultaneously regulates membrane transport, protein degradation, and gene expression. To address this, we examined whether Ca^2+^ signals imposed via XXM2.0 alter auxin-induced gene expression in the absence of the exogenous auxin treatment. To this end, three-day-old Arabidopsis seedlings expressing XXM2.0, GtACR1, and the controls grown in an RL background were exposed to 1s BL every 30s. Roots challenged by BL stimulation were harvested after 1 h, 8 h, 32 h for RNA-seq. Global transcriptomic analyses uncovered XXM2.0 as a potent tool to modulate gene expression. Principal component analysis (PCA) revealed that the GtACR1 plants (function as depolarization control of XXM2.0) clustered together at all time points; in contrast, XXM2.0 samples diverged markedly upon BL stimulation (Fig. S6). Among the differentially expressed genes (DEGs), we observed the induction of classical Ca^2+^ signaling markers such as *CML24/TCH2*, whose transcripts are known to rise following mechanical or cold-induced Ca^2+^ transients (*43, 44*). In addition, key Ca^2+^ decoders, including *CPKs*, *CAMTAs*, and *CAMs* and were also upregulated (Fig. S7), indicating that optogenetic Ca^2+^ input activates endogenous gene expression programs associated with Ca^2+^ signaling.

Time-series clustering of XXM2.0-regulated DEGs revealed five distinct clusters. Although one cluster of Ca^2+^-responsive genes was enriched in defense-related transcriptional regulators, auxin-responsive genes grouped into distinct clusters with separate temporal dynamics. Specifically, auxin-associated DEGs in clusters 1, 4 and 5 included canonical transcriptional targets such as *AUX2-11, AUX2-27, IAA30*, and *AXR5*, with peak expression at 1 h, followed by a decline at 8 and 32 h (Fig. 4H). Notedly, no auxin biosynthesis genes were found in the DEGs. Gene Ontology (GO) analysis of these clusters revealed significant enrichment for terms including ‘response to auxin’, ‘regulation of root morphogenesis’ (Fig. 4H-I and Fig. S6).

To identify the Ca^2+^ sensitive tissues and cells in the seedling roots, we mapped 45 auxin-associated genes from clusters 1, 4, and 5 onto the Arabidopsis root atlas (Fig. S8A). Of these, 36 genes were highly correlated with specific tissues. Among the root tissue types, the protoxylem was the most strongly enriched cell type, expressing 11 of the 36 genes (Fig. S8B). Several of these - including *IAA30*, *BG1*, *PATL5*, and *SAUR19* - were robustly downregulated in protoxylem cells by up to ∼2-fold at 8 h. In the columella, which exhibited pronounced *DR5v2*::n3GFP suppression and contributes to root cap signaling, genes such as *ACC4*, *SAUR10*, and *SAUR50* were notably reduced (∼1.5–2× decrease) in XXM2.0 roots (Fig. S8C). Similarly, cortical and epidermal cells showed downregulation of *ATXTH19*, *TET10*, and *AXR5*, genes involved in cell wall remodeling and auxin-regulated elongation (Fig. S8C). Importantly, gene expression in the quiescent center and distal stem cell niche remained largely unchanged (Fig. S8C), suggesting that auxin suppression is not uniform but targeted to specific differentiation and elongation domains. Together, these data reveal that optogenetically imposed Ca^2+^ signals via XXM2.0 reprogram auxin-responsive gene expression in a spatially and temporally defined manner, demonstrating that consecutive cytosolic Ca^2+^ transients are sufficient to reshape key auxin transcriptional networks in Arabidopsis roots.

## Optogenetic Ca^2+^ stimulation arrests root growth in a reversible manner

Because environmental stress-induced Ca^2+^ signals were proposed to modulate growth (*45*), we next asked whether endogenous Ca^2+^ transients elicited by natural stimuli similarly affect root elongation. We adopted a microfluidic root imaging platform to simultaneously monitor cytosolic Ca^2+^ dynamics and root growth rates in real time (*46*). GCaMP3-expressing seedlings were grown within the microchip and exposed to 5 mM glutamate, a wound-associated signaling molecule known to trigger cytosolic Ca^2+^ transients in roots (Fig. 5A-B). Glutamate application rapidly induced a pronounced cytosolic Ca^2+^ signal, which was accompanied by an immediate reduction in root growth rate. Importantly, repetitive glutamate perfusion pulses generated repeated Ca^2+^ transients that synchronized with repeated decreases in growth rate. To further test whether this phenomenon extends to abiotic stress, we tested the response of cold-induced Ca^2+^ signaling (Fig. 5C-D). A rapid temperature shift (22 °C to 14 °C) provoked a strong cytosolic Ca^2+^ transient and a marked reduction in root growth rate. Notably, upon returning seedlings to 22°C, root growth recovered to pre-treatment levels, indicating that this response is reversible and dynamically regulated. Together, these findings show that stress-induced Ca^2+^ transients are tightly associated with rapid and reversible suppression of root growth, as has also been shown for other stimuli (*47*).

**Fig. 5.**
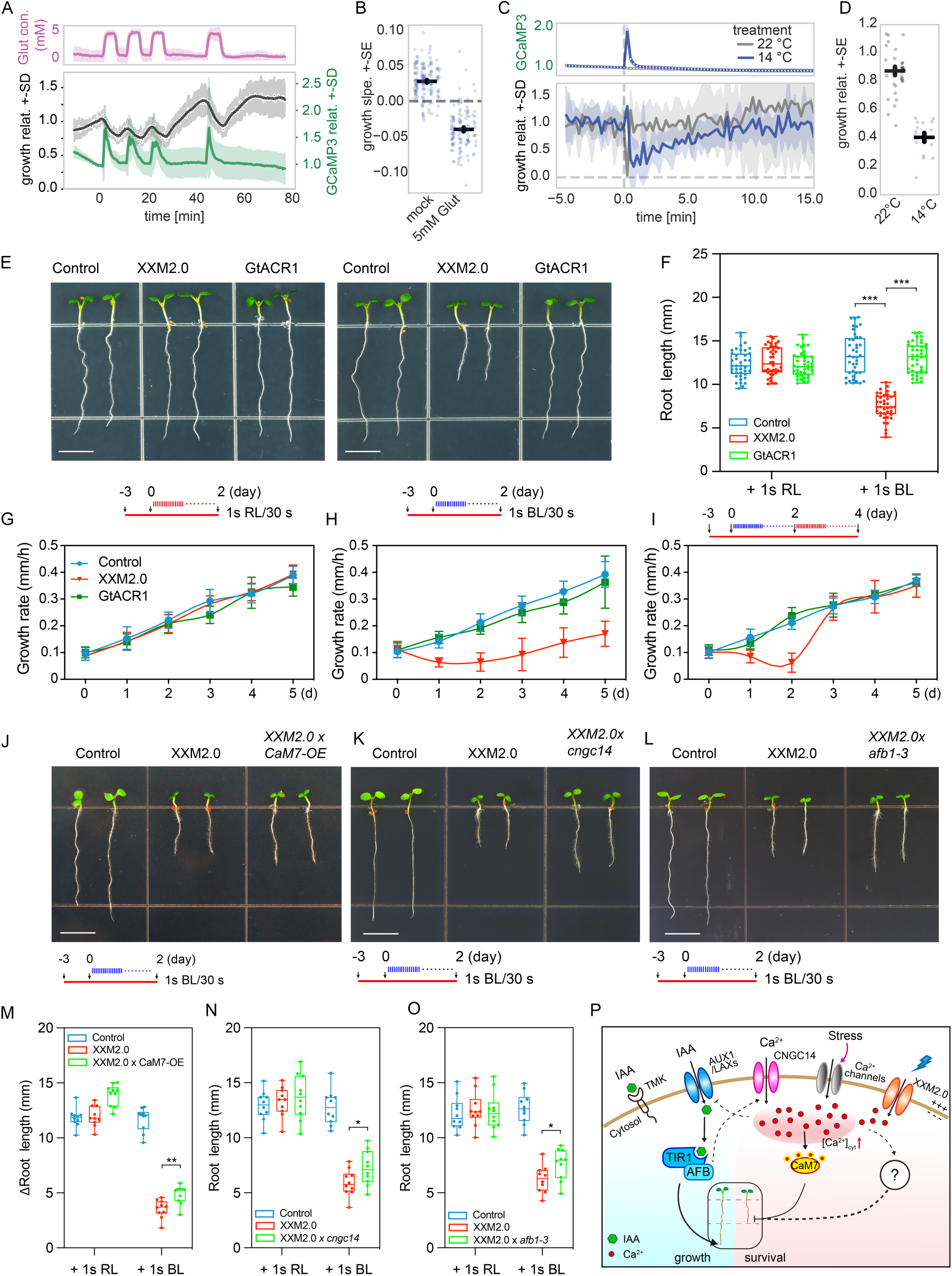
Ca^2+^ signals inhibit root growth in a reversible manner. (A) Simultaneous monitoring of root growth rate and cytosolic Ca^2+^ dynamics in Arabidopsis roots expressing GCaMP3 during repetitive local application of 5 mM glutamate (5 s). Upper panel: glutamate treatment regime. Lower panel: relative root growth rate (black) and GCaMP3 fluorescence (green). Shaded areas indicate ± SD. (B) Quantification of root growth slope following mock or 5 mM glutamate treatment. Each dot represents one biological replicate; black bars indicate mean ± SE. (C) Relative root growth rate and GCaMP3 fluorescence in Arabidopsis roots exposed to cold treatment (14 °C) or maintained at 22 °C. Dashed line indicates the onset of temperature treatment. Shaded areas indicate ± SD. Inset: representative GCaMP3 fluorescence traces under 22 °C and 14 °C conditions. (D) Quantification of relative root growth under 22 °C and 14 °C conditions. Each dot represents one biological replicate; black bars indicate mean ± SE. (**E**) Root length phenotypes of 5-day-old transgenic *Arabidopsis* lines (Control, XXM2.0, and GtACR1). Seedlings were grown under continuous red LED illumination (650 nm, 100 µE) for 3 days, followed by stimulation with RL or BL pulses (1 s every 30 s) for an additional 2 days. Scale bars = 5 mm. (**F**) Quantification of primary root length from (E). Data are mean ± SD (*n* = 40 biological replicates). *** *p* < 0.001. (**G-H**) Root growth rate of Control, XXM2.0, and GtACR1 seedlings under RL (E) or BL (F) pulsed stimulation for 5 days period. Data are mean ± SD (*n* = 10). (**I**) Root growth rate of Control, XXM2.0, and GtACR1 seedlings stimulated with a 2 d BL + 2 d RL pulsed light. Data are mean ± SD (*n* = 10). **(J-L**) Root length phenotypes of 5-day-old transgenic Arabidopsis lines Control, XXM2.0, and XXM2.0 x CaM7-OE (J), XXM x *cngc14* (K), XXM x *afb1*(L). Seedlings were grown under continuous red LED illumination (650 nm, 100 μE) for 3 days, and followed by stimulation with RL or BL pulses (1 s every 30 s) for an additional 2 days. Scale bars = 5 mm. **(M-O)** Quantification of primary root length in Control, XXM2.0, and XXM2.0 x *CaM7-OE* or *cngc14*, *afb1-3* seedlings after 2 days of RL or BL pulse light treatment. Root length was calculated as the final primary root length minus the primary root length before RL/BL pulses treatment for XXM2.0 x *CaM7-OE*. (n = 10 biological replicates). * *p* < 0.05, ** *p* < 0.01. Scale bars = 5 mm. (**P**) Working model for Ca^2+^-gated early auxin signaling in roots.

These observations prompted us to ask whether repeated Ca^2+^ inputs are not only sufficient to transiently inhibit root elongation, but can also drive longer-term developmental changes in root architecture. To test this hypothesis, XXM2.0, GtACR1, and control seedlings were grown under RL for 3 days, followed by 2 additional days of RL or BL stimulation. After 2 days of RL pulse stimulation, the phenotype of roots from GtACR1, XXM2.0, and the control showed similar primary root lengths (Fig. 5E-F). This finding was confirmed by growth rate measurements (Fig. 5G-H). Following BL pulse treatment, the situation did not change fundamentally when comparing the control and GtACR1 roots. The XXM2.0 roots, however, after 2 days exhibited a marked growth arrest with primary root length reduced to approximately 50% of that in the control and GtACR1 plants (Fig. 5E-F). Daily growth rate analysis shows that BL-stimulated XXM2.0 roots expanded only ∼40% the rate of the controls and GtACR1 across multiple days, with the maximal divergence observed on day one (Fig. 5H). Next, we asked if the apparent BL-induced arrest of XXM2.0 root growth is reversible. To this end, 2 days BL challenged plants were switched to 2 days of RL stimulation. Following this protocol, XXM2.0 roots rapidly resumed growth, with growth rates comparable to control and GtACR1 plants within 24 h (Fig. 5I). Root elongation increased by 33% during recovery, indicating that BL/XXM2.0-induced growth arrest is reversible (Fig. S12A-B).

To analyze the restricted growth of BL-stimulated XXM2.0 roots in more detail, we microscopically inspected the root meristem, elongation zone, and differentiation zone. For better visualization of cell borders, cell walls were stained with Calcofluor White. Under RL conditions, root growth zone architecture and cell length distributions were nearly identical among the controls and XXM2.0 and GtACR1 lines, with cortex cells in each zone reaching up to ∼280 µm, 420 µm, and 11,000 µm in length, respectively (Fig. S9A-B, E and H). In contrast, we found that after 2 days of BL-stimulation, the meristematic, elongation, and differentiation zones of XXM2.0 were reduced by 32%, 65%, and 76%, respectively, compared to both the control and GtACR1 (Fig. S9A, C, F, I). Concerning the cell number in the three root zones, we found that under BL conditions, XXM2.0 roots were reduced by about 32% and 45% the number in the meristem and differentiation zones compared to the control and GtACR1 plants, and that of the elongation zones were less reduced (Fig. S9). Seedlings switched from BL to RL, in the newly developed root region, cell lengths and numbers in the meristematic and elongation zones recovered to levels comparable to those of the control, and the differentiation zone also reinitiated growth (Fig. S9A, D, G, J). These observations suggest that persistent optogenetically-induced Ca^2+^ stimulations reversibly repress cell division and elongation, leading to inhibited root growth.

To further assess whether the developmental defects of XXM2.0 roots result from changes in endogenous auxin signalling, we examined additional auxin-dependent growth phenotypes. Under BL stimulation, XXM2.0 seedlings showed strongly reduced lateral root number and density, as well as impaired gravitropic bending, whereas control and GtACR1 roots remained largely unaffected (Fig. S10). These classical auxin-related phenotypes are also consistent with Ca^2+^ dependent modulation of endogenous auxin signalling in roots.

To test whether the proposed CaM7–CNGC14 module also contributes to the developmental output of optogenetically imposed Ca^2+^ signals, we analysed root growth in XXM2.0 × CaM7-OE seedlings. Under RL conditions, XXM2.0 × CaM7-OE seedlings displayed longer primary roots than both control and XXM2.0 lines (Fig. S12C), consistent with the elongated root phenotype of CaM7-OE and *cngc14* plants. Following BL stimulation, root growth remained strongly inhibited in XXM2.0 × CaM7-OE seedlings, but the reduction in root elongation was significantly weaker than in XXM2.0 plants, resulting in greater growth increments (Δroot length; Fig. 5J, 5M and Fig. S12). These results indicate that XXM-imposed Ca^2+^ signals have a weaker inhibitory effect on root growth in the CaM7-OE background. Likewise, disruption of CNGC14 or AFB1 partially alleviated BL-induced root growth inhibition, indicating that endogenous auxin signalling is associated with Ca^2+^-imposed growth arrest (Fig. 5K-O and Fig. S12).

## Discussion

### Ca^2+^ acts as a stop signal to modulate early auxin signalling during environmental stress

In response to biotic and abiotic changes in the environment, plants must decide when to grow and when to pause. Auxin signaling plays a central role in regulating root growth and architecture, positioning it as a potentially critical decision-making hub under environmental stress. However, the signals that dynamically modulate early auxin signalling remained largely unknown due to the lack of non-invasive tools for cytoplasmic Ca^2+^ stimulation.

Here, we imposed defined cytosolic Ca^2+^ signatures via XXM2.0 optogenetically and documented that Ca^2+^ acts as a central gating signal for early auxin signalling in Arabidopsis roots. By dissecting early Ca^2+^–electrical auxin signalling events, our findings demonstrate that preceding cytosolic Ca^2+^ signals dynamically tune the strength of early auxin signalling. Short XXM2.0-triggered Ca^2+^ transients, described previously as all-or-none “sparks” (*26*), progressively attenuated auxin-induced membrane depolarization and cytosolic Ca^2+^ transients (Fig. 2), demonstrating that preceding cytosolic Ca^2+^ signals dynamically tune the strength of early auxin signalling. Beyond these rapid signalling events, the same Ca^2+^ signals also suppressed auxin-responsive transcription and altered auxin-dependent development (Fig. 4 and Fig. 5), revealing that Ca^2+^ regulates auxin responses across multiple levels of the auxin response. Importantly, these effects were dose-dependent and reversible, underscoring the dynamic and tunable nature of Ca^2+^-dependent regulation of auxin signalling.

Crucially, this Ca^2+^-dependent gating is not limited to optogenetic inputs. Physiological stimuli such as cold and glutamate, and even auxin itself, known to provoke cytosolic Ca^2+^ transients, triggered comparable reductions in auxin responsiveness (Fig. 3). Notably, XXM-induced Ca^2+^ priming also attenuated glutamate- and cold-triggered Ca^2+^ transients (Fig. S11), indicating that elevated cytosolic Ca^2+^ can transiently reduce the general Ca^2+^ responsiveness of root cells. Thus, Ca^2+^-dependent regulation of early auxin signalling may represent one line of a broader feedback mechanism controlling stimulus-evoked Ca^2+^ excitability. Together, these findings support a model in which cytosolic Ca^2+^ elevations act as a transient “stop signal”, temporarily uncoupling growth from hormonal inputs during stress episodes and enabling roots to prioritize survival over elongation. Importantly, the impact of Ca^2+^ signalling on auxin responses are not uniform across the root. While auxin is classically viewed as an inhibitor of elongation at high concentrations (*48–50*), under basal conditions it also promotes meristem activity and cell expansion (*48–50*). Thus, when auxin responses are broadly attenuated by Ca^2+^, both inhibitory and promotive signaling outputs are suppressed. DR5 reporter signals and transcriptomic analyses indicate that auxin responsiveness is especially reduced in elongation- and differentiation-related tissues, such as the cortex, protoxylem, epidermis, and columella—while the stem cell niche remains relatively unaffected (Fig. S8). This spatially restricted decline in auxin-responsive outputs likely underlies the observed phenotype: smaller elongation zones, shorter cells, and reversible inhibition of primary root growth (Fig. 5). In this way, Ca^2+^ gating functions as a stress-responsive brake, allowing plants to transiently halt growth in specific tissues while maintaining developmental competence.

### How does Ca^2+^ regulate early auxin signaling at the molecular level?

At first glance, the observed attenuation of auxin responsiveness may appear counterintuitive, as auxin signalling itself depends on cytosolic Ca^2+^ transients. Our data instead support a model in which Ca^2+^ acts as a transient negative feedback signal that dampens subsequent auxin responsiveness (Fig. 2 and Fig. 3). AUX1-mediated auxin uptake and AFB1-dependent signalling are known to activate CNGC14-dependent Ca^2+^ influx during early auxin responses (*13, 15, 18, 19*), raising the possibility that Ca^2+^ elevations feed back onto this pathway to limit further activation. In this context, CNGC14 represents an attractive candidate node for feedback regulation (*13*), particularly because the Ca^2+^ sensor CaM7 has previously been shown to inhibit CNGC14 activity in a heterologous system (*40*). In line with this, CaM7-overexpression plants phenocopied *cngc14* mutants, displaying strongly impaired auxin-induced membrane depolarization and cytosolic Ca^2+^ signals, as well as elongated primary roots (Fig. 3 and Fig. S3). The ability of CaM7-overexpression alone to suppress auxin responsiveness further suggests that elevated CaM7 reduces the sensitivity of this signalling branch, consistent with the weaker effect of optogenetically imposed Ca^2+^ signals on root growth in CaM7 × XXM seedlings (Fig. 5J). Likewise, disruption of either CNGC14 or AFB1 partially alleviated XXM-induced root growth inhibition (Fig. 5K–O), indicating that the early auxin signalling pathway represents a crucial, but not the sole, target of the Ca^2+^-dependent feedback. Together, these findings support a model in which a preceding Ca^2+^ transient engages CaM7 to transiently suppress CNGC14-mediated auxin signalling, thereby limiting subsequent auxin responsiveness (Fig. 5P). This points to a tightly coupled feedback mechanism at the plasma membrane, through which preceding Ca^2+^ signals can rapidly tune auxin responsiveness at an early signalling step, providing a plausible means to transiently adjust growth decisions following stress-associated Ca^2+^ elevations. How auxin perception engages the CaM7–CNGC14 module, and how this feedback is executed at the molecular level, now emerge as key questions for future investigation.

Beyond this fast response, our data indicate that Ca^2+^ stimulation can also reshape auxin outputs over longer timescales without altering IAA homeostasis or PIN-dependent auxin transport (Fig. S4 and S5). These observations suggest that major changes in auxin biosynthesis or polar transport are less likely to underlie the observed effects. Instead, our results using the DR5 transcriptional auxin reporter point towards suppression of canonical transcriptional auxin signalling itself (Fig. 4). In line with this, transcriptome analysis revealed widespread downregulation of classical auxin-responsive genes, including *AUX/IAAs* and *SAURs*, following prolonged XXM2.0 activation. Because these genes are regulated by canonical signaling components such as AUX1 and TIR1/AFBs, CNGC14, Aux/IAAs and ARFs (*41*), attenuation of early auxin responsiveness through the CaM7–CNGC14 branch may progressively reshape downstream transcriptional outputs. However, whether this mechanism alone is sufficient to drive the observed transcriptional changes remains an open question.

In addition to auxin-responsive genes, our RNA-seq data show an induction of *CMLs, CPKs*, and *CAMTAs* (Fig. S7), all known Ca^2+^-regulated proteins with potential to interact with auxin-related targets. For instance, CaM/CMLs can modulate scaffolds such as IQD proteins, influencing the activity of key kinase in auxin biology, PINOID, or affect early auxin response genes (*51–53*), suggesting that Ca^2+^ decoding mechanisms could modulate auxin responses at multiple levels, although effects on local auxin distribution remain to be resolved. CAMTAs have been implicated in transcriptional reprogramming under stress, and may contribute to Ca^2+^-dependent auxin gating through modulation of gene expression (*54*).

### How do distinct Ca^2+^ signatures regulate auxin signalling?

An outstanding question is how distinct Ca^2+^ signatures are translated into specific auxin outputs. Optogenetics provides a unique opportunity to dissect how defined Ca^2+^ signatures regulate auxin signalling, a question that has previously been difficult to address experimentally. Within the parameter range tested here, different XXM-imposed Ca^2+^ patterns converged on a similar attenuation of auxin responsiveness (Fig. S2). This may reflect rapid saturation of the underlying feedback mechanism or sustained regulation of key auxin signalling components by repeated Ca^2+^ transients. Consistent with this idea, our previous guard cell study showed that different Ca^2+^ stimulation patterns primarily affected the kinetics rather than the final extent of stomatal closure (*26*). Exploring a broader range of Ca^2+^ signatures should provide new insights into how Ca^2+^ dynamics regulate auxin signalling.

### Ca^2+^ signalling beyond early auxin signaling

Although our data identify early auxin signalling as a major target of Ca^2+^ priming in roots, Ca^2+^ is well known to function as a central signalling hub across diverse physiological pathways. Consistent with this broader role, prolonged XXM2.0 activation induced transcriptional signatures associated not only with auxin regulation, but also with salicylic acid and immune-related pathways (Fig. S6). These observations suggest that defined Ca^2+^ stimulation can engage additional downstream stress tolerance programs beyond auxin signalling, particularly during prolonged stimulation.

At the same time, the rapid and reversible effects observed here, for example, the growth inhibition, occurring within minutes after XXM2.0 activation and similarly reproduced by endogenous Ca^2+^ transients triggered by cold, glutamate, and auxin (Fig. 5). Our findings indicate that the modulation of early auxin signalling pathway contributes to the growth inhibition, although additional Ca^2+^-regulated pathways also contribute to longer-term effects. Natural stimuli likely provide additional layers of information through receptor activation, membrane voltage changes, ROS, peptide signalling, and metabolic cues, which may shape downstream signalling specificity. In this context, the synthetic biology optogenetic approach XXM2.0 provides a tractable framework to dissect which aspects of complex environmental responses can be attributed to Ca^2+^ signalling itself.

In conclusion, our findings establish cytosolic Ca^2+^ as a dynamic regulator of early auxin signalling. By acting as a reversible gating signal, Ca^2+^ enables plants to transiently decouple auxin-mediated growth signaling from stress-induced survival programs, facilitating adaptive plasticity. Optogenetic control of Ca^2+^ via XXM2.0 offers a powerful tool to dissect this hormone– second messenger crosstalk with unprecedented temporal resolution. Future studies integrating optogenetics with mutants of Ca^2+^ decoders such as CaMs, CAMTAs, CIPKs and CPKs will further illuminate how plants prioritize growth versus survival in fluctuating environments.

## Materials and Methods

### Plasmid construction and plant transformation

The XXM2.0 coding sequence was amplified from genomic DNA of *Nicotiana tabacum* lines expressing XXM2.0 (*26, 27*) using gene-specific primers (Table S1). The resulting cDNA was inserted into the pCAMBIA3300 binary vector under the control of the UBQ10 promoter using USER cloning (*55*). This construct also included the *MbDio* gene (marine bacterial *β-carotene 15,15′-dioxygenase*) to enable retinal biosynthesis (*29*), as well as a USER cassette upstream of an eYFP reporter for expression verification (*28*). A version of the plasmid lacking eYFP was also generated. Binary constructs were introduced into *Arabidopsis thaliana* Col-0 plants via the *Agrobacterium tumefaciens* strain GV3101 using the floral dip method (*56*). Transgenic seedlings were selected on soil using BASTA (0.2% Glufosinate Ammonium). For Ca^2+^ imaging, the genetically encoded indicator R-GECO1 was amplified from the R-GECO1-mTurquoise plasmid (*20, 57*), and two R-GECO1 units were tandemly linked using the peptide linker GLNLSGG, as previously described (*58*). The construct was also driven by the UBQ10 promoter and cloned into pCAMBIA3300. *CaM7* was amplified from Arabidopsis cDNA using USER-compatible primers and cloned into a pCAMBIA2300-based USER cassette, generating a C-terminal mCherry fusion construct under the control of the UBQ10 promoter. Primer sequences used for cloning are listed in Table S1.

### Plant materials and growth conditions

All *Arabidopsis* lines used in this study were in the Col-0 background. The XXM2.0 and GtACR1 lines (lacking eYFP) was crossed with *DR5v2*::n3GFP-*Tcsn*::ntdTomato (*59*). GtACR1 and control (expressing a retinal synthesis gene and a YFP) lines were generated previously (*28*). Plants were grown in sterilized soil under climate-controlled conditions with red LED illumination (650 nm, 100 µmol·m^-2^·s^-1^). The photoperiod was set to 16 h light / 8 h dark, with day/night temperatures of 22°C and 18°C, respectively, and relative humidity maintained at 60%.

For root phenotyping, *Arabidopsis* seeds were surface-sterilized with 6% (v/v) NaClO, containing 0.05% (v/v) Triton X-100 for 10 min and washed three to six times with sterilized distilled water. Sterilized seeds were sown on modified ½ MS medium containing (1% sucrose and 1.7% agar at pH 5.8 with TRIS) and stratified at 4°C in the dark for 3 days to synchronize germination. Plates were then placed vertically in a growth chamber at 22°C under continuous red LED illumination (650 nm, 100 µmol·m^-2^·s^-1^). Three-day-old seedlings with comparable root size were stimulated with red light (RL) or blue light (BL) pulses (duration = 1 s, time interval = 30 s, intensity = 100 µmol·m^-2^·s^-1^) for 2 days, unless otherwise specified in the figure legends.

### Electrophysiological recordings on root cells

Electrophysiological recordings were performed on cortical cells in the root meristematic zone. Four- to six-day-old seedlings were gently fixed onto a Petri dish using medical adhesive (Medical Adhesive B; Aromando, Germany) and incubated in a bath solution containing 1 mM KCl, 1 mM CaCl2, and 5 mM MES/BTP (pH 6.0), unless otherwise specified. The Petri dish was then placed on the stage of an upright microscope (Axioskop 2FS, Zeiss, Germany) and kept there for four hours prior to measurements. Root cells were impaled using sharp microelectrodes pulled from borosilicate glass capillaries (inner/outer diameter = 0.56/1.0 mm; Hilgenberg, Germany) with a horizontal laser puller (P-2000, Sutter Instruments, CA, USA). Electrodes were filled with 300 mM KCl and had a tip resistance of 150-180 MΩ. A reference electrode, filled with 300 mM KCl sealed with 2% agarose, was used as a reference. The electrodes were mounted into a holder attached to a piezo-driven micromanipulator (MM3A, Kleindiek, Reutlingen, Germany), which was used to impale root cells under microscopic guidance. The microelectrodes were connected via Ag/AgCl half-cells to headstages with an input impedance of 100 GΩ. These were further connected to a custom-built amplifier (Ulliclamp01). Electrical signals were low-pass filtered at 0.5 kHz using a dual low-pass Bessel filter (LPF-202A; Warner Instruments Corp., USA) and recorded at 1 kHz using a USB-6002 data acquisition interface (National Instruments, USA) operated by WinWCP or WinEDR software (*60*).

### Local IAA application

Glass capillaries were pulled using a horizontal puller and mounted on a micromanipulator (Sensapex). For local applications, pipette tips were broken to a diameter of approximately 10 µm and positioned ∼200 µm from the root tip. A pressure of 2 bar was applied for 5 s using a Picospritzer II microinjection system (npi, Germany), corresponding to an estimated ejection volume of 200 nL. Prior to IAA application, the impaled cells were allowed to stabilize for several minutes. A 10 µM IAA solution (dissolved in bath medium) was then applied for 5 s. This was followed by a 10-minute wash period to allow the membrane potential to return to baseline.

Data were processed using custom Python scripts, and statistical analyses were performed in GraphPad Prism 8.0. Raw recordings were filtered with a Gaussian filter (sigma = 1000) and down-sampled to 10 Hz. The 10-s period preceding the IAA puff was used to define baseline values; the mean of this baseline was subtracted from the rest of the recording to determine the absolute IAA-evoked depolarization. For dose–response analysis, the depolarization amplitude from the second IAA application was normalized to the amplitude of the first IAA response. The normality and homogeneity of variance for each dataset were assessed, followed by statistical testing.

### Auxin reporter assay

For auxin assays, transgenic plants were generated by crossing the auxin reporter line *DR5v2::n3GFP* with XXM2.0 or GtACR1 (without eYFP). The applied BL regime was first confirmed to induce sustained, repetitive cytosolic Ca^2+^ transients in XXM2.0 roots throughout the recording period (Fig. S13). Following light treatment, seedlings were fixed in 4% (w/v) paraformaldehyde at room temperature, washed twice with 1 × PBS for 5 minutes each, and kept in the dark until imaging. Confocal images of root tips were acquired, and fluorescence was quantified using ImageJ Software (imagej.nih.gov/ij). Regions of interest (ROIs) were drawn around the root tip in each image, and corrected total cell fluorescence (CTCF = Integrated density – (Area × mean background) was calculated for each ROI.

### Measurement of root length and growth rate

Root length was measured from the digital photographs of the plates using RootDetection (www.labutils.de) and ImageJ (http://rsb.info.nih.gov/ij/). Each root length was recorded as the final after completion of the specific light stimulation. Root growth rate was calculated by measuring the daily increase in root length and dividing this value by 24 h to obtain the growth rate per hour.

### Measurement of cell length and cell number

Root cell length and number were measured by staining seedlings with Calcofluor White (Sigma-Aldrich, 18909). Seedlings were fixed in 4% (w/v) paraformaldehyde in 1× PBS for 1 hour at room temperature under vacuum. Then washed twice with 1× PBS for 1 min each. Samples were cleared overnight in ClearSee solution containing 10% (w/v) xylitol powder, 15% (w/v) sodium deoxycholate, and 25% (w/v) urea with gentle agitation, as previously described (*61*). Next, seedlings were incubated in 0.1% Calcofluor White in Clearsee solution for 30 min, rinsed for 30 min in Clearsee solution. Mount on slides in Clearsee for imaging.

The meristematic zone was defined from the quiescent center to the last cell that had not doubled in size, the elongation zone from the first cell that doubled in size to the last cell before root hair bulges, and the differentiation zone from the first cell below a trichoblast bulge to the root-shoot junction. All measurements were performed on all individual cells of a consecutive cortex cell file using ImageJ.

### Ca^2+^ imaging

Ca^2+^ imaging was performed as previously described (*28*). Arabidopsis seedlings expressing the red fluorescent Ca^2+^ indicator R-GECO1 were excited using an LED illumination system (pE-4000; CoolLED, Andover, UK) at 580 nm. Emission signals were filtered through a dichroic mirror with a 590 nm cut-off wavelength (FF593 BrightLine; Semrock) and a bandpass filter centered at 628/40 nm (BrightLine; Semrock). The dichroic mirror and emission filter were mounted in the filter wheels of a CARV II spinning disk confocal unit (CrestOptics). Images were acquired using a scientific CMOS camera (Prime BSI; Teledyne Photometrics) controlled by VisiView software (Visitron) and processed using the Fiji software package. IAA was locally applied to the root surface in the meristematic zone, and R-GECO1 signals were recorded from the same region.

### PIN immuno-labeling

The whole-mount immune-labelling protocol (*62*) was applied with some modifications. Seedlings were fixed in 4% paraformaldehyde in PBS for 30min in the vacuum desiccator, and then washed in PBS as previously described for cell length and cell number measurements. For automated immuno-labeling by Intavis InSitu Pro robot we followed a previously published protocol (*63*). We incubated 3 times for 15 min in PBS + 0.1% Triton X-100 at RT, followed by 3 times 15 min in ddH2O + 0.1% Triton X-100. Cell-wall digestion was performed for 30 min with 2% driselase in PBS at 37 °C. Samples were washed 3 times for 15 min in PBS + 0.1% Triton X-100 at RT. Permeabilization was performed twice for 30 min with 10% DMSO and 3% Igepal CA-630 in PBS at RT, followed by 3 times washing for 15 min in PBS + 0.1% Triton X-100 at RT. Blocking was done for 1 h with 2% BSA in PBS. Samples were incubated for 4 h at 37 °C in blocking solution containing custom antibodies raised in rabbit against PIN1 (*64*) or PIN2 (*65*). Samples were washed 3 times for 15 min at RT in PBS + 0.1% Triton X-100. Secondary anti-rabbit Cy3 conjugated antibody (Abcam ab6939, dilution 1:600) in blocking solution was applied at 37 °C for 4 h. Samples were washed 3 times for 15 min in PBS + 0.1% Triton X-100 at RT and then 3 times 15 min in ddH2O at RT. The next day, seedlings were mounted in 90% Glycerol in PBS, pH 8,0, with 25mg/ml DABCO. The cover glass was framed by transparent nail polish. Imaging was done on Axio Imager.Z2 equipped by LSM800 and Plan-Apochromat 40x/1.2 water objective.

### Confocal laser scanning microscopy

Microscopy images were performed using a confocal laser scanning microscope (Leica SP5, Leica Microsystems, Germany), controlled by Leica LAS AF (v2.7.3.9723), and Nikon ECLIPSE Ti2, controlled by NIS-Elements 5.4. For subcellular localization of eYFP in root cells, yellow fluorescence was imaged with a 25× HCX IRAPO 925/0.95 W dipping objective, excited at 496 nm and detected at 520-580 nm. Calcofluor White staining was imaged with the same objective, using 405 nm excitation and detected at 425-475 nm. For the auxin reporter line DR5v2::n3GFP, green fluorescence in the root cells was captured with a 40× /1.10 CORR CS dipping objective, and 40× /0.95 OFN25 DIC N2, excited at 488 nm and detected at 505-530 nm.

### Transcriptome analysis

Illumina sequencing reads (150bp, paired end) were generated by Novogene using an Illumina NovaSeq 6000. Quality control and read mapping was done using “amalgkit integrate”, “amalgkit getfastq” and “amalgkit merge” from the AMALGKIT (version 0.12.7, https://github.com/kfuku52/amalgkit) toolkit. Reads were mapped against TAIR10 *Arabidopsis thaliana* CDS sequences obtained from The Arabidopsis Information Resource (https://www.arabidopsis.org). Differential gene expression analysis was then performed in R (version 4.4.1) using with DESeq2 package (version 1.4.4) (*66*). For each genotype and each timepoint, a comparison was made between blue light treated samples (Condition A) versus the red light treated control samples (Condition B). For a gene to be considered differentially expressed, it had to pass the following filters: padj < 0.05, log2FC > 1 and for downregulated genes: padj < 0.05, log2FC < -1. 2517 DEGs identified as unique to the XXM genotype across all timepoints of all genotypes were then selected for the cluster analysis. Gene counts in the form of TPM were log2 transformed prior to clustering. Clustering was done with the clusterProfiler Package (version 4.12.6) (*67*) with a k = 5. Z-score normalized gene expression of clustered genes was plotted using pheatmap (version 1.0.12). For tissue-specific expression analysis in the root, six high-quality datasets (Table S2) were retrieved from scPlantDB (*68*). Tissue specificity was assessed using the τ index, and average gene expression was calculated for each tissue and gene.

### Quantification of auxin metabolites

Auxin metabolites were quantified as described previously (*69*). Approximately 10 mg fresh weight of root samples was extracted with 1 mL 50 mmol/L phosphate buffer (pH 7.0) containing 0.1% sodium diethyldithiocarbamate and stable isotope-labelled internal standards. Aliquots of 200 µL were either acidified to pH 2.7 and purified by in-tip µSPE, or derivatized with cysteamine prior to acidification and µSPE for IPyA determination. Eluates were evaporated, reconstituted in 10% aqueous methanol, and analyzed using an Agilent 1260 Infinity II HPLC with a Kinetex C18 column (50 × 2.1 mm, 1.7 µm) coupled to an Agilent 6495 Triple Quadrupole mass spectrometer.

### Measurement of lateral root number and lateral root density

Transgenic Arabidopsis thaliana seedlings (Control, XXM2.0, and GtACR1) were vertically grown on ½ MS medium under continuous red LED illumination (650 nm, 100 μE) for 3 days, followed by treatment with RL or BL pulses (1 s every 30 s) for an additional 5 days. Lateral root formation was assessed in 8-day-old seedlings by counting sites of lateral root initiation or visible lateral root primordia using a standard light microscope (Leica EZ4HD, Leica, Germany). Images were captured with a Nikon BM-10 and analyzed with ImageJ. Lateral root density was calculated as the number of lateral roots per unit length of the primary root.

### Measurement of root gravitropism

Transgenic Arabidopsis seedlings (control, XXM2.0, and GtACR1) were vertically grown on ½ MS medium under continuous red LED illumination (650 nm, 100 μE) for 3 days. And then the plates were rotated 90° counterclockwise, and the seedlings were then maintained vertically under RL or BL pulses (1 s every 30 s) for an additional 4 h. Root curvature was recorded after treatment using a standard light microscope (Leica EZ4HD, Leica, Germany) and camera (Nikon BM-10), and the curvature was quantified using ImageJ, with the original vertical root growth direction defined as the baseline.

### Quantification of auxin-induced DR5v2::n3GFP responses

To examine the effect of auxin on reporter expression, transgenic Arabidopsis seedlings expressing DR5v2::n3GFP were vertically grown on ½ MS medium for 72 h. Seedlings were then spray-treated with 1 μM IAA prepared from a DMSO stock solution. Mock-treated seedlings were sprayed with the corresponding solvent control containing the same final concentration of DMSO. Root GFP fluorescence was then monitored at 1, 8, 24, and 48 h after treatment. Root tips were imaged at each time point using a confocal laser scanning microscope (Leica SP5, Leica Microsystems, Germany), controlled by Leica LAS AF (v2.7.3.9723). Images were acquired with a 40×/1.10 CORR CS dipping objective, with GFP excited at 488 nm and detected at 505-530 nm. For fluorescence quantification, the root tips were selected as the region of interest (ROI), and GFP intensity was measured in ImageJ as the mean gray value. Data represent n = 8 roots per group and are presented in arbitrary units (a.u.).

### Cold-induced root inhibition assay

For cold treatment, four-day-old GCaMP3 seedlings were mounted externally beneath a bottom-glass chamber (Lab-Tek™ II). The seedlings were embedded in 1/2 MS medium supplemented with 1% sucrose, pH 5.8 (AM+), and secured in place with a coverslip. The chamber was filled with 1 ml of room temperature water and the treatment was done either by adding 1ml of either room temperature or ice-cooled water. Imaging was performed using a Nikon Ti2-E inverted wide-field microscope equipped with a Plan Fluor 10× objective (NA = 0.3). In a separate benchtop experiment under identical environmental conditions, the temporal temperature profile of the system was characterized using an infrared thermometer (ABOHU, G1).

### Glutamate treatment in the RootChip system

Four-day-old GCaMP3 seedlings were treated using the RootChip microfluidics system as previously described (*47*). In brief, a single-layer polydimethylsiloxane (PDMS) chip containing two inlets and a branching channel network feeding a 1 cm wide, 110 μm deep observation chamber for roots was used. The device was fabricated by mixing RTV615 PDMS in a 5:1 ratio, degassing, pouring the mixture over a wafer mold, and baking overnight at 80°C. Following excision, 0.5 mm inlet holes were made using a puncher (RapidCore, 0.5 mm), and 1/32’’ ID tubing was connected using 22G pins. Seedlings were placed into the open observation chamber containing AM+ medium and sealed with a coverslip; the glass–PDMS assembly was clamped together in a screw-tightened holder to form a microscope insert. Flow in the chip was maintained at 20 μL/min using two NE-300 syringe pumps for glutamate treatment or mock treatment, respectively. Solution exchange was monitored using TRITC-dextran dye (Sigma-Aldrich, T1162). After addition of 5 mM L-glutamic acid (Sigma-Aldrich, 49449) to the AM+ medium, the pH was adjusted to match the mock treatment using KOH (Merck, 105021), and osmolality in the mock solution was balanced with sorbitol (Sigma-Aldrich, S1876; approximately 10 mM), according to measurements using an Osmomat 3000 (Gonotec). Imaging was performed on a vertically oriented Axio Observer (Zeiss) wide-field microscope equipped with a Plan-Apochromat 10× objective (NA = 0.45).

### Root growth image analysis

Image analysis was performed as previously described (*47*). In brief, using ImageJ (v1.54p), the approximate root midline was manually tracked from the differentiation zone to the root tip by placing ordered points across multiple time frames. A custom Python script was then used for downstream analysis. The script interpolated the coordinates in XY space and time, establishing regularly spaced square regions along the root line. Subpixel registration on bright-field images (image_registration v0.2.10, chi2_shift) was used to quantify overall root shifts (growth, bending, and stage drift) within these regions. Root elongation was calculated by summing distance changes between adjacent regions while correcting for bending and stage drift. The average fluorescence intensity of the GCaMP channel was also quantified within each region during the analysis pipeline.

## Acknowledgments

We would like to thank Dirk Becker (University of Wuerzburg) for his help in making the optogenetic constructs and for his comments on the manuscript. We also appreciate Rob Roelfsema (University of Wuerzburg) for his feedback on the manuscript. We sincerely thank the anonymous reviewers for the thoughtful and inspiring suggestions that helped broaden the discussion and identify important future research directions.

## Funding

National Key R&D Program of China, 2025YFA0923400 (RH)

National Natural Science Foundation of China (NSFC), W2531024 (RH); 32271501(NL)

NSFC-Excellent Young Scientists Fund Program (overseas) (SH)

NSFC-RFIS, ZB2025101410702 (RH)

Guangdong Pearl River Talents Program, 2023ZT10N033

China Scholarship Council no. 202306510063 (HS)

The European Research Council grant 101142681 CYNIPS (JF)

Austrian Science Fund grant FWF; P 37051-B (JF)

German Research Foundation grant CRC 1644 (512328399; 512328399) (KK, ZH)

ERC Synergy project STARMORPH, reg. no. 101166880 (AP, ON)

## Author contributions

Conceptualization: SH, RH

Methodology: SH, HS, AB, MF, MR, AP, ZH, KK, NO, JF, NL

Investigation: SH, HS, AB, MF, MR, AP, ZH

Visualization: RH, SH, JF, MG, KK, NO, HS, AB, MF, MR, AP, ZH, NL

Funding acquisition: RH, SH, KK, JF

Project administration: RH, SH

Supervision: RH, SH, KK, JF

Writing – original draft: RH, SH

Writing – review & editing: RH, SH, JF, MG, KK, NO, HS, AB, MF, MR, AP, ZH

## Competing interests

Authors declare that they have no competing interests.

## Data and materials availability

All data are available in the main text or the supplementary materials.

## Notes

### Competing Interest Statement

The authors have declared no competing interest.

### Summary of Updates

We used more precise words to describe auxin responsiveness and also added a summarized figure.

## References

1. W. Lin et al., TMK-based cell-surface auxin signalling activates cell-wall acidification. Nature 599, 278–282 (2021).

2. L. Li et al., Cell surface and intracellular auxin signalling for H+ fluxes in root growth. Nature 599, 273–277 (2021).

3. H. Chen et al., TIR1-produced cAMP as a second messenger in transcriptional auxin signalling. Nature 640, 1011–1016 (2025).

4. M. Ke et al., Salicylic acid regulates PIN2 auxin transporter hyperclustering and root gravitropic growth via Remorin-dependent lipid nanodomain organisation in Arabidopsis thaliana. New Phytol 229, 963–978 (2021).

5. D. Das, K. R. St Onge, L. A. Voesenek, R. Pierik, R. Sasidharan, Ethylene- and Shade-Induced Hypocotyl Elongation Share Transcriptome Patterns and Functional Regulators. Plant Physiol 172, 718–733 (2016).

6. J.-E. Park et al., GH3-mediated Auxin Homeostasis Links Growth Regulation with Stress Adaptation Response in Arabidopsis. Journal of Biological Chemistry 282, 10036–10046 (2007).

7. H. Suda et al., Calcium dynamics during trap closure visualized in transgenic Venus flytrap. Nat Plants 6, 1219–1224 (2020).

8. S. Scherzer et al., A unique inventory of ion transporters poises the Venus flytrap to fast-propagating action potentials and calcium waves. Current Biology, 32, 4255–4263. (2022).

9. M. Toyota et al., Glutamate triggers long-distance, calcium-based plant defense signaling. Science 361, 1112–1115 (2018).

10. F. Wu et al., Hydrogen peroxide sensor HPCA1 is an LRR receptor kinase in Arabidopsis. Nature, 578, 577–581 (2020).

11. F. Yuan et al., OSCA1 mediates osmotic-stress-evoked Ca^2+^ increases vital for osmosensing in Arabidopsis. Nature 514, 367–371 (2014).

12. Z. Jiang et al., Plant cell-surface GIPC sphingolipids sense salt to trigger Ca2+ influx. Nature, 572, 341–346 (2019).

13. J. Dindas et al., AUX1-mediated root hair auxin influx governs SCF(TIR1/AFB)-type Ca2+ signaling. Nat Commun 9, 1174 (2018).

14. R. Wang et al., Auxin analog-induced Ca2+ signaling is independent of inhibition of endosomal aggregation in Arabidopsis roots. J Exp Bot 73, 2308–2319 (2022).

15. H. W. Shih, C. L. DePew, N. D. Miller, G. B. Monshausen, The Cyclic Nucleotide-Gated Channel CNGC14 Regulates Root Gravitropism in Arabidopsis thaliana. Curr Biol 25, 3119–3125 (2015).

16. N. B. C. Serre et al., The AUX1-AFB1-CNGC14 module establishes a longitudinal root surface pH profile. Elife 12, e85193 (2023).

17. S. Vanneste, Y. Pei, J. Friml, Mechanisms of auxin action in plant growth and development. Nat Rev Mol Cell Biol 26, 648–666 (2025).

18. M. Fendrych et al., Rapid and reversible root growth inhibition by TIR1 auxin signalling. Nat Plants 4, 453–459 (2018).

19. N. B. C. Serre et al., AFB1 controls rapid auxin signalling through membrane depolarization in Arabidopsis thaliana root. Nature Plants 7, 1229–1238 (2021).

20. S. Huang et al., Calcium signals in guard cells enhance the efficiency by which abscisic acid triggers stomatal closure. New Phytol 224, 177–187 (2019).

21. G. J. Allen, K. Kuchitsu, S. P. Chu, Y. Murata, J. I. Schroeder, Arabidopsis *abi1-1* and *abi2-1* phosphatase mutations reduce abscisic acid-induced cytoplasmic calcium rises in guard cells. Plant Cell 11, 1785–1798 (1999).

22. G. J. Allen et al., A defined range of guard cell calcium oscillation parameters encodes stomatal movements. Nature 411, 1053–1057 (2001).

23. M. R. McAinsh, A. Webb, J. E. Taylor, A. M. Hetherington, Stimulus-Induced Oscillations in Guard Cell Cytosolic Free Calcium. Plant Cell 7, 1207–1219 (1995).

24. A. Reyer et al., Channelrhodopsin-mediated optogenetics highlights a central role of depolarization-dependent plant proton pumps. Proc Natl Acad Sci U S A, 117, 20920–20925 (2020).

25. S. Huang et al., Optogenetic control of the guard cell membrane potential and stomatal movement by the light-gated anion channel GtACR1. Science Advances 7, eabg4619 (2021).

26. S. Huang, M. R. G. Roelfsema, M. Gilliham, A. M. Hetherington, R. Hedrich, Guard cells count the number of unitary cytosolic Ca2+ signals to regulate stomatal dynamics. Current Biology, 34, 5409–5416. (2024).

27. M. Ding et al., Probing plant signal processing optogenetically by two channelrhodopsins. Nature, 633, 872–877 (2024).

28. S. Huang, L. Shen, M. R. G. Roelfsema, D. Becker, R. Hedrich, Light-gated channelrhodopsin sparks proton-induced calcium release in guard cells. Science 382, 1314–1318 (2023).

29. Y. Zhou et al., Optogenetic control of plant growth by a microbial rhodopsin. Nature Plants 7, 144–151 (2021).

30. J. J. Jones, S. Huang, R. Hedrich, C. M. Geilfus, M. R. G. Roelfsema, The green light gap: a window of opportunity for optogenetic control of stomatal movement. New Phytol, 236, 1237–1244 (2022).

31. R. Hedrich, M. Gilliham, Light-activated channelrhodopsins: a revolutionary toolkit for the remote control of plant signalling. New Phytol 245, 982–988 (2025).

32. W. G. Choi, M. Toyota, S. H. Kim, R. Hilleary, S. Gilroy, Salt stress-induced Ca2+ waves are associated with rapid, long-distance root-to-shoot signaling in plants. Proc Natl Acad Sci U S A 111, 6497–6502 (2014).

33. C. Allan et al., Observing root growth and signalling responses to stress gradients and pathogens using the bi-directional dual-flow RootChip. Lab on a chip 24, 5360–5373 (2024).

34. J. Dindas et al., Pitfalls in auxin pharmacology. New Phytol, 227, 286–292. (2020).

35. S. Huang, R. Hedrich, Trigger hair thermoreceptors provide for heat-induced calcium-electrical excitability in Venus flytrap. Current Biology 33, 3962–3968.e3962 (2023).

36. A. Carpaneto et al., Cold transiently activates calcium-permeable channels in Arabidopsis mesophyll cells. Plant Physiology 143, 487–494 (2007).

37. M. R. Knight, H. Knight, Low-temperature perception leading to gene expression and cold tolerance in higher plants. New Phytologist 195, 737–751 (2012).

38. Y. Yu et al., ABLs and TMKs are co-receptors for extracellular auxin. Cell 186, 5457–5471.e5417 (2023).

39. J. Friml et al., ABP1-TMK auxin perception for global phosphorylation and auxin canalization. Nature 609, 575–581 (2022).

40. Q. Zeb et al., The interaction of CaM7 and CNGC14 regulates root hair growth in Arabidopsis. Journal of Integrative Plant Biology 62, 887–896 (2020).

41. C. Luschnig, J. Friml, Over 25 years of decrypting PIN-mediated plant development. Nat Commun 15, 9904 (2024).

42. C.-Y. Liao et al., Reporters for sensitive and quantitative measurement of auxin response. Nature Methods 12, 207–210 (2015).

43. M. Matsumura et al., Mechanosensory trichome cells evoke a mechanical stimuli-induced immune response in Arabidopsis thaliana. Nat Commun 13, 1216 (2022).

44. D. H. Polisensky, J. Braam, Cold-shock regulation of the Arabidopsis TCH genes and the effects of modulating intracellular calcium levels. Plant Physiol 111, 1271–1279 (1996).

45. S. Luan, Calcium signaling in plants: Universal and unique paradigms. Cell 189, 1001–1023 (2026).

46. I. Kulich et al., Calcium-triggered apoplastic ROS bursts balance gravity and mechanical signals for soil navigation. Science 392, 296–300 (2026).

47. M. Randuch et al., Cytosolic Ca2+ as a universal signal for rapid root growth regulation. bioRxiv, 2025.2010.2017.683082 (2025).

48. M. L. Evans, R. E. Cleland, The action of auxin on plant cell elongation. Critical Reviews in Plant Sciences 2, 317–365 (1985).

49. R. Cleland, The dosage-response curve for auxin-induced cell elongation: A reevaluation. Planta 104, 1–9 (1972).

50. L. Taiz, I. M. Møller, A. Murphy, E. Zeiger, Plant physiology and development. (2023).

51. S. Zhang et al., The calcium signaling module CaM-IQM destabilizes IAA-ARF interaction to regulate callus and lateral root formation. Proc Natl Acad Sci U S A 119, e2202669119 (2022).

52. R. Benjamins, C. S. Ampudia, P. J. Hooykaas, R. Offringa, PINOID-mediated signaling involves calcium-binding proteins. Plant Physiol 132, 1623–1630 (2003).

53. H. J. Teresinski et al., Arabidopsis calmodulin-like proteins CML13 and CML14 interact with proteins that have IQ domains. Plant, Cell & Environment 46, 2470–2491 (2023).

54. Y. Galon et al., Calmodulin-binding transcription activator 1 mediates auxin signaling and responds to stresses in Arabidopsis. Planta 232, 165–178 (2010).

55. H. H. Nour-Eldin, B. G. Hansen, M. H. Norholm, J. K. Jensen, B. A. Halkier, Advancing uracil-excision based cloning towards an ideal technique for cloning PCR fragments. Nucleic acids research 34, e122 (2006).

56. X. Zhang, R. Henriques, S. S. Lin, Q. W. Niu, N. H. Chua, Agrobacterium-mediated transformation of Arabidopsis thaliana using the floral dip method. Nature protocols 1, 641–646 (2006).

57. R. Waadt, M. Krebs, J. Kudla, K. Schumacher, Multiparameter imaging of calcium and abscisic acid and high-resolution quantitative calcium measurements using R-GECO1-mTurquoise in Arabidopsis. New Phytologist 216, 303–320 (2017).

58. K. Li et al., An optimized genetically encoded dual reporter for simultaneous ratio imaging of Ca2+ - and H+ reveals new insights into ion signaling in plants. New Phytol, 230, 2292–2310 (2021).

59. C. Y. Liao et al., Reporters for sensitive and quantitative measurement of auxin response. Nat Methods 12, 207–210, 202 p following 210 (2015).

60. J. Dempster, A new version of the Strathclyde Electrophysiology software package running within the Microsoft Windows environment. Journal of Physiology 504, P57–P57 (1997).

61. D. Kurihara, Y. Mizuta, Y. Sato, T. Higashiyama, ClearSee: a rapid optical clearing reagent for whole-plant fluorescence imaging. Development 142, 4168–4179 (2015).

62. M. Karampelias, R. Tejos, J. Friml, S. Vanneste, Optimized Whole-Mount In Situ Immunolocalization for Arabidopsis thaliana Root Meristems and Lateral Root Primordia. Methods Mol Biol 1761, 131–143 (2018).

63. M. Sauer, T. Paciorek, E. Benková, J. Friml, Immunocytochemical techniques for whole-mount in situ protein localization in plants. Nature protocols 1, 98–103 (2006).

64. T. Paciorek et al., Auxin inhibits endocytosis and promotes its own efflux from cells. Nature 435, 1251–1256 (2005).

65. L. Abas et al., Intracellular trafficking and proteolysis of the Arabidopsis auxin-efflux facilitator PIN2 are involved in root gravitropism. Nature cell biology 8, 249–256 (2006).

66. M. I. Love, W. Huber, S. Anders, Moderated estimation of fold change and dispersion for RNA-seq data with DESeq2. Genome biology 15, 550 (2014).

67. S. Xu et al., Using clusterProfiler to characterize multiomics data. Nature protocols 19, 3292–3320 (2024).

68. Z. He et al., scPlantDB: a comprehensive database for exploring cell types and markers of plant cell atlases. Nucleic acids research 52, D1629–D1638 (2023).

69. A. Pencík et al., Ultra-rapid auxin metabolite profiling for high-throughput mutant screening in Arabidopsis. J Exp Bot 69, 2569–2579 (2018).

