## Supplementary material for "Ca^2+^ signature gates early auxin signaling in Arabidopsis roots": suppl. info

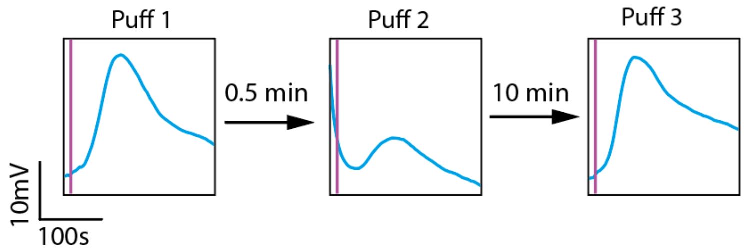


**Fig. S1. IAA sensitivity recovers after termination of XXM2.0-induced $Ca^{2+}$ signals.**
Representative membrane potential recordings from a XXM2.0-expressing root cell during sequential IAA stimulations. Puff 1: first local IAA application induces a typical depolarization. Following a blue light train (λ = 470 nm, 100 µE, 20 pulses of 1 s at 30 s intervals), Puff 2 shows a markedly reduced depolarization. After 10 min recovery, Puff 3 elicits a restored membrane depolarization.

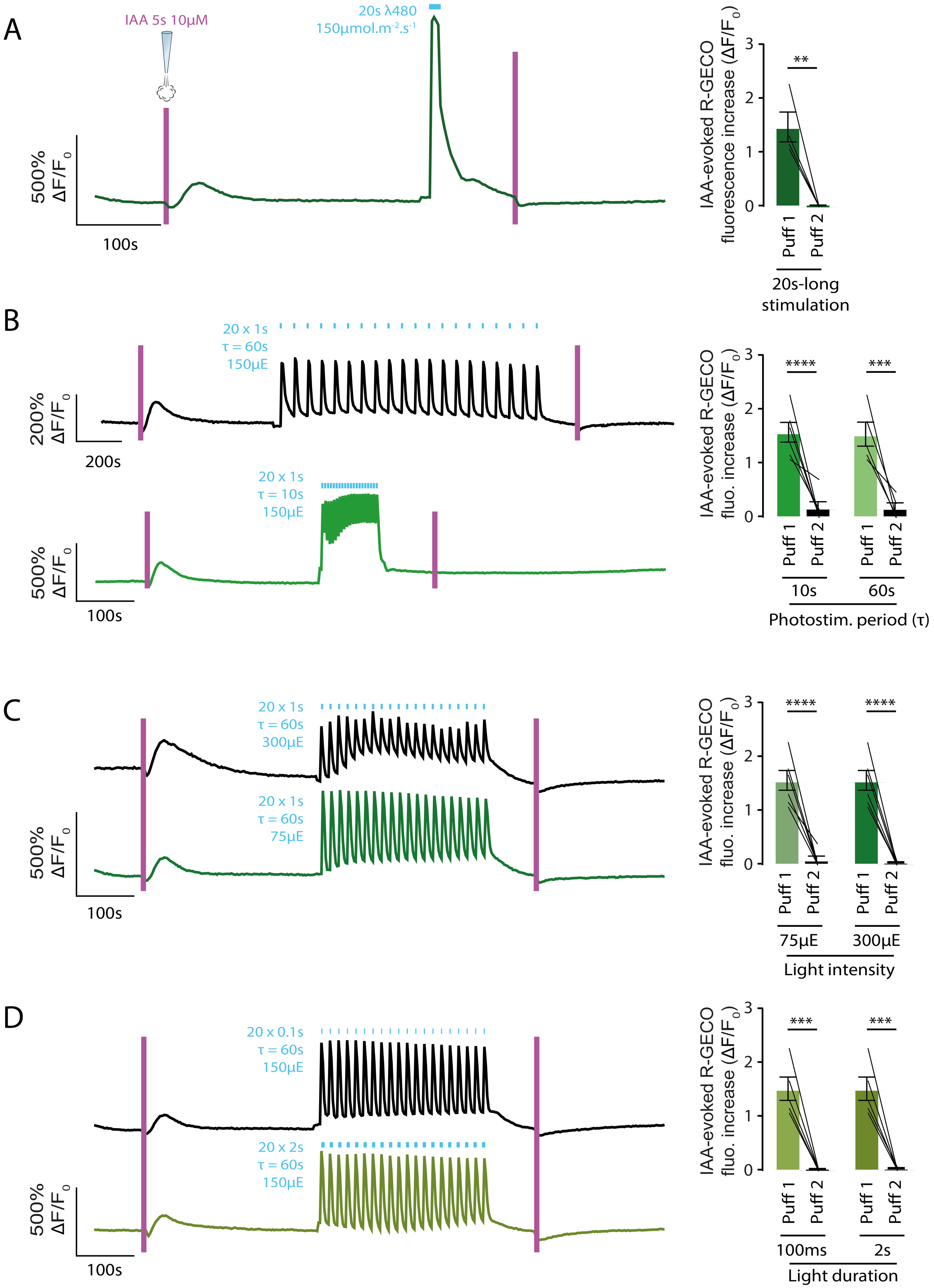
A
IAA 5s 10µM
20s λ480
150µmol.m-2.s-1
500% ΔF/F0
100s
IAA-evoked R-GECO fluorescence increase (ΔF/F0)
**
Puff 1
Puff 2
20s-long stimulation
B
20 x 1s
τ = 60s
150µE
200% ΔF/F0
200s
20 x 1s
τ = 10s
150µE
500% ΔF/F0
100s
IAA-evoked R-GECO fluo. increase (ΔF/F0)
****
***
10s
60s
Photostim. period (τ)
C
20 x 1s
τ = 60s
300µE
20 x 1s
τ = 60s
75µE
500% ΔF/F0
100s
****
****
75µE
300µE
Light intensity
D
20 x 0.1s
τ = 60s
150µE
20 x 2s
τ = 60s
150µE
500% ΔF/F0
100s
***
***
100ms
2s
Light duration

1018
1019

**Fig. S2 Optogenetically induced $Ca^{2+}$ transients suppress subsequent auxin-induced $Ca^{2+}$ responses.**

(A) Representative R-GECO1 fluorescence trace showing $Ca^{2+}$ responses evoked by two sequential 10 μM IAA (5 s) applications in Arabidopsis root meristematic cells before and after XXM2.0 activation by 20 s BL ($\lambda$ = 480 nm, 150 μmol $m^{-2}$ $s^{-1}$). Right panel: quantification of IAA-evoked R-GECO1 fluorescence increase ($\Delta F/F_0$). (B) Representative R-GECO1 fluorescence traces and quantification of IAA-evoked $Ca^{2+}$ responses following XXM2.0 activation using 20 × 1 s BL pulses delivered at 10 s or 60 s intervals ($\tau$) at 150 μmol $m^{-2}$ $s^{-1}$. (C) Representative R-GECO1 fluorescence traces and quantification of IAA-evoked $Ca^{2+}$ responses following XXM2.0 activation using 20 × 1 s BL pulses ($\tau$ = 60 s) at 75 or 300 μmol $m^{-2}$ $s^{-1}$. (D) Representative R-GECO1 fluorescence traces and quantification of IAA-evoked $Ca^{2+}$ responses following XXM2.0 activation using 20 BL pulses of either 100 ms or 2 s duration ($\tau$ = 60 s, 150 μmol $m^{-2}$ $s^{-1}$). Bars represent mean ± SE. Each line connects paired biological replicates. $^{**}p < 0.01$, $^{***}p < 0.001$, $^{****}p < 0.0001$.

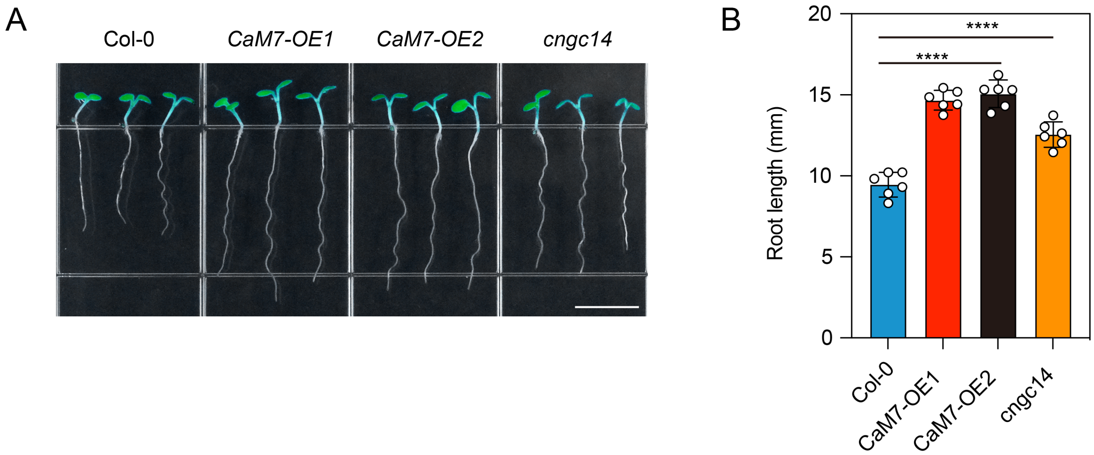


**Fig. S3. CaM7-overexpression plants exhibit elongated primary root phenotypes similar to *cngc14*.**

(A) Representative images of 5-day-old Arabidopsis seedlings of Col-0, CaM7-OE#1, CaM7-OE#2 (independent UBQ10::CaM7-mCherry overexpression lines), and *cngc14*. Seedlings were grown vertically under white LED illumination. Scale bar = 5 mm. (B) Quantification of primary root length in Col-0, CaM7-OE#1, CaM7-OE#2, and *cngc14* seedlings. Bars represent SD (n = 6 biological replicates). ****$p < 0.0001$.

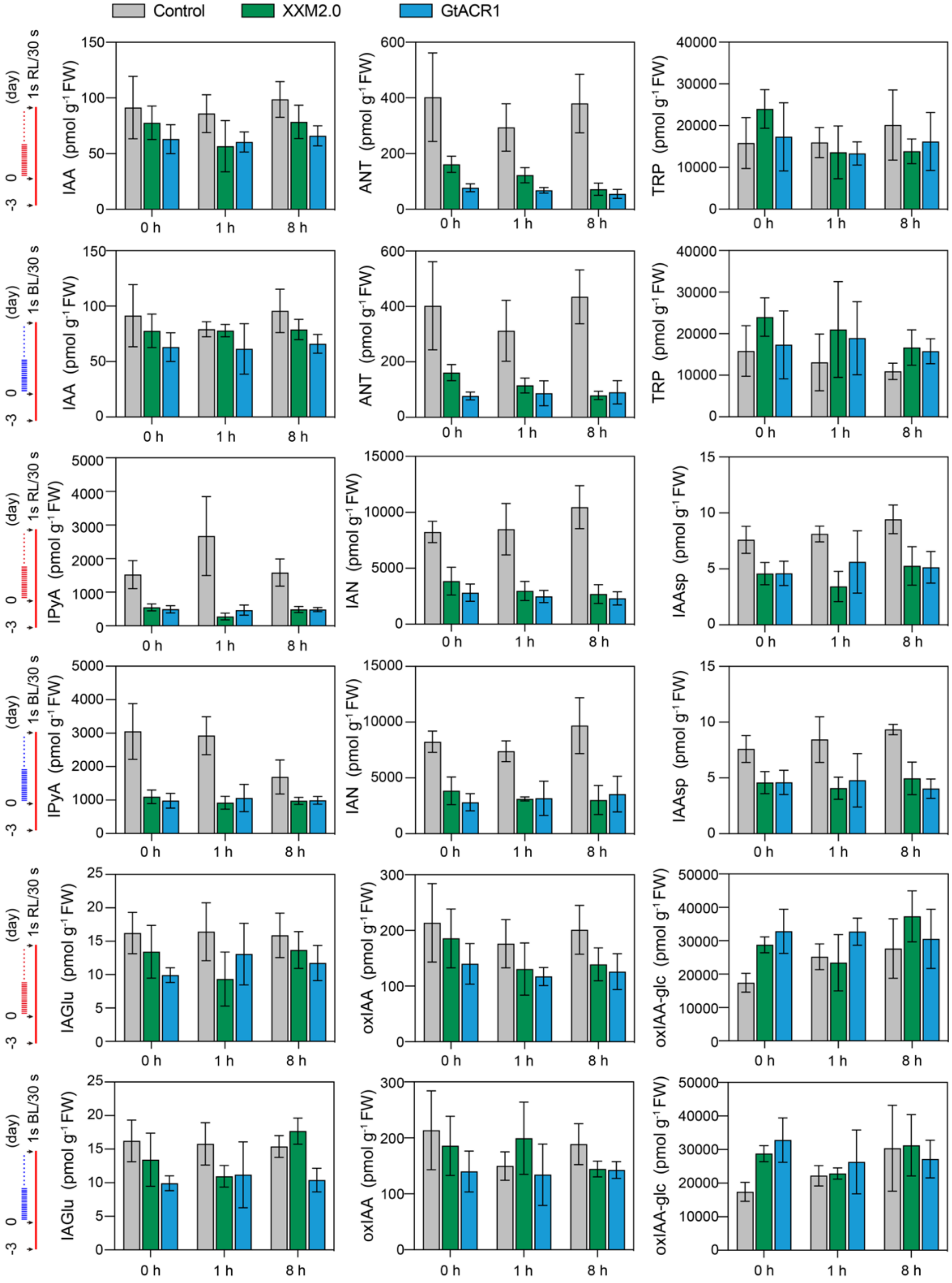
Control
XXM2.0
GtACR1
(day)
1s RL/30 s
1s BL/30 s
-3
0
IAA (pmol g-1 FW)
ANT (pmol g-1 FW)
TRP (pmol g-1 FW)
IPyA (pmol g-1 FW)
IAN (pmol g-1 FW)
IAAsp (pmol g-1 FW)
IAGlu (pmol g-1 FW)
oxIAA (pmol g-1 FW)
oxIAA-glc (pmol g-1 FW)
0 h
1 h
8 h

**Fig. S4. Quantification of IAA biosynthesis and metabolic intermediates in Arabidopsis roots.**

Levels of IAA and associated biosynthetic and catabolic intermediates were measured in roots of control, XXM2.0, and GtACR1 plants after 0, 1, and 8 h of light treatment. Plants were grown for 3 days under RL, followed by 1-s BL pulses every 30 s for the indicated durations. Metabolite quantification was performed via HPLC. No significant differences in IAA content or related metabolite levels were detected between XXM2.0 and GtACR1 samples at any time point. Error bars indicate standard error (n = 5).

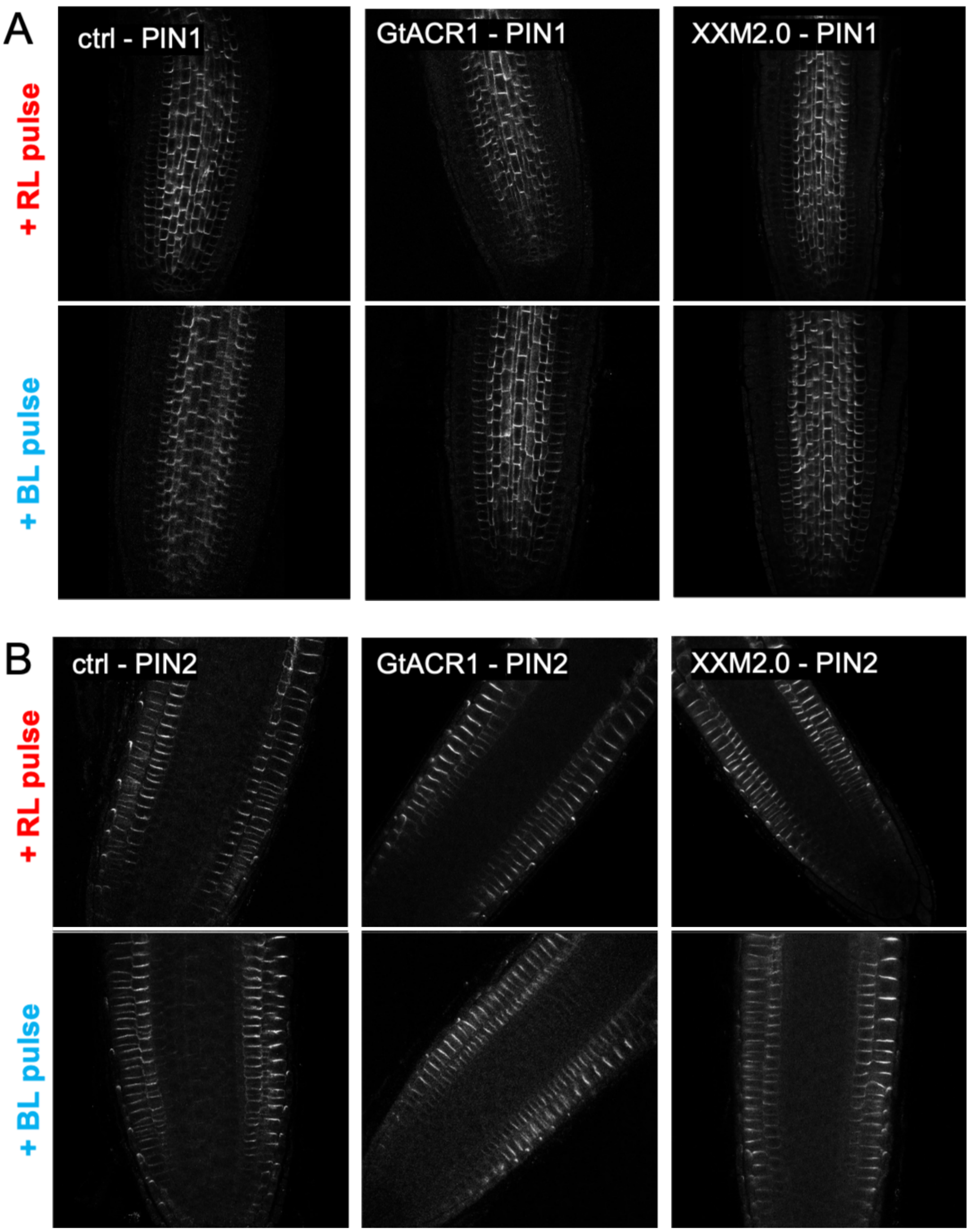


**Fig. S5. PIN1 and 2 immunostaining in roots of controls, XXM2.0, and GtACR1.**
Immunostaining of PIN1 (A) and PIN2 (B) in Arabidopsis seedling roots expressing either XXM2.0, GtACR1, or controls. Seedlings were grown for 2 days under continuous RL or BL stimulation (1 s pulse every 30 s, 100 µE). Roots were fixed and immunolabeled with anti-PIN1 or anti-PIN2 antibodies. For each genotype and treatment, at least 10 roots were imaged from two independent biological replicates. Shown are representative confocal images with contrast adjustments applied for clarity. No differences in PIN1 or PIN2 localization patterns were observed across conditions.

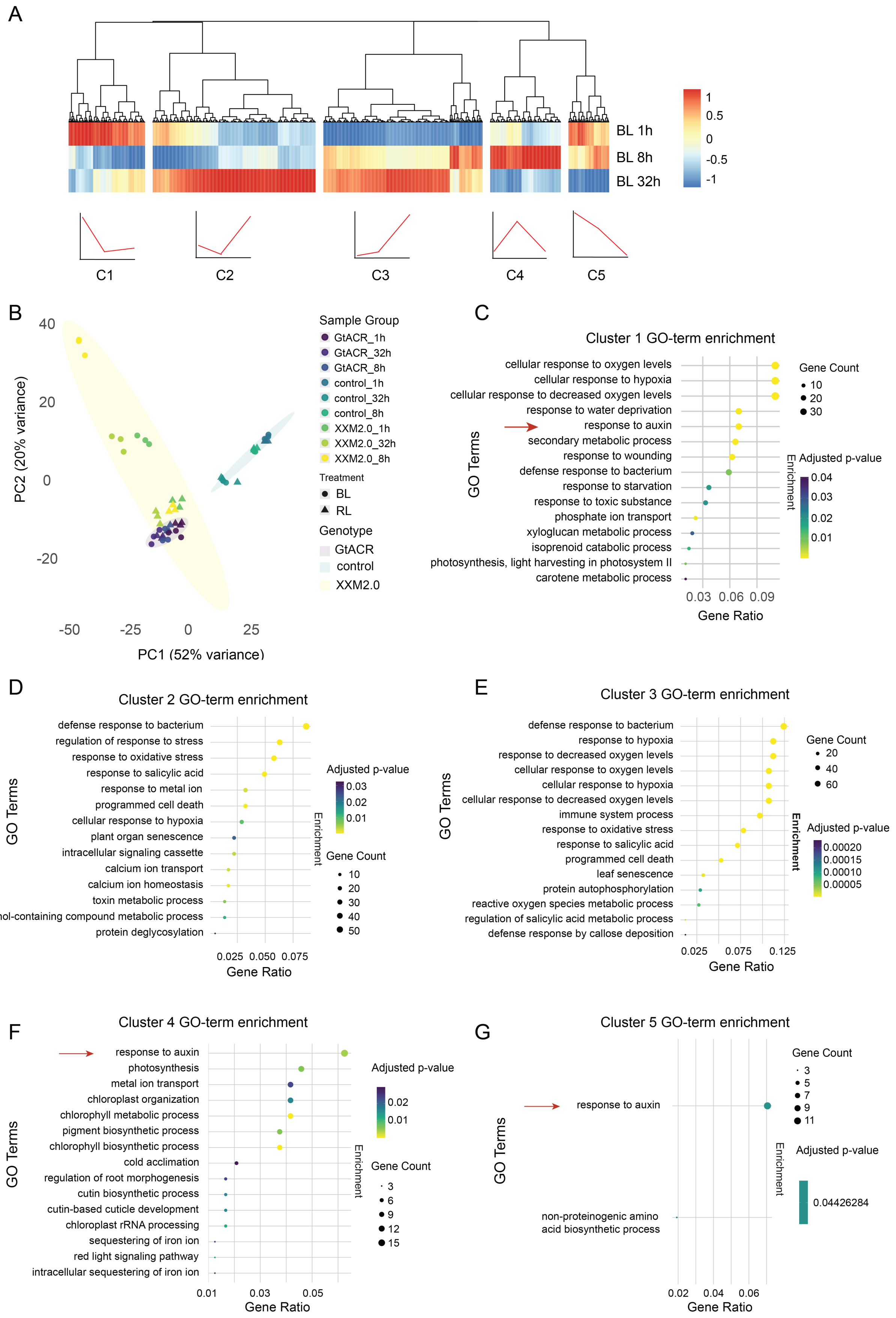
A
BL 1h
BL 8h
BL 32h
1
0.5
0
-0.5
-1
C1
C2
C3
C4
C5
B
PC2 (20% variance)
PC1 (52% variance)
Sample Group
GtACR_1h
GtACR_32h
GtACR_8h
control_1h
control_32h
control_8h
XXM2.0_1h
XXM2.0_32h
XXM2.0_8h
Treatment
BL
RL
Genotype
GtACR
control
XXM2.0
C
Cluster 1 GO-term enrichment
cellular response to oxygen levels
cellular response to hypoxia
cellular response to decreased oxygen levels
response to water deprivation
response to auxin
secondary metabolic process
response to wounding
defense response to bacterium
response to starvation
response to toxic substance
phosphate ion transport
xyloglucan metabolic process
isoprenoid catabolic process
photosynthesis, light harvesting in photosystem II
carotene metabolic process
GO Terms
Gene Ratio
Gene Count
Adjusted p-value
Enrichment
D
Cluster 2 GO-term enrichment
defense response to bacterium
regulation of response to stress
response to oxidative stress
response to salicylic acid
response to metal ion
programmed cell death
cellular response to hypoxia
plant organ senescence
intracellular signaling cassette
calcium ion transport
calcium ion homeostasis
toxin metabolic process
phenol-containing compound metabolic process
protein deglycosylation
E
Cluster 3 GO-term enrichment
defense response to bacterium
response to hypoxia
response to decreased oxygen levels
cellular response to oxygen levels
cellular response to hypoxia
cellular response to decreased oxygen levels
immune system process
response to oxidative stress
response to salicylic acid
programmed cell death
leaf senescence
protein autophosphorylation
reactive oxygen species metabolic process
regulation of salicylic acid metabolic process
defense response by callose deposition
F
Cluster 4 GO-term enrichment
response to auxin
photosynthesis
metal ion transport
chloroplast organization
chlorophyll metabolic process
pigment biosynthetic process
chlorophyll biosynthetic process
cold acclimation
regulation of root morphogenesis
cutin biosynthetic process
cutin-based cuticle development
chloroplast rRNA processing
sequestering of iron ion
red light signaling pathway
intracellular sequestering of iron ion
G
Cluster 5 GO-term enrichment
response to auxin
non-proteinogenic amino acid biosynthetic process
0.04426284

**Fig. S6. Differential gene expression and functional enrichment analysis of $Ca^{2+}$-stimulated Arabidopsis roots.**

(**A**) Hierarchical clustering of differentially expressed genes (DEGs) across XXM2.0 roots at 0 h, 1 h, and 32 h after BL stimulation. Five distinct gene expression clusters (C1–C5) were identified based on temporal expression dynamics. (**B**) Principal component analysis (PCA) of transcriptomes reveals clear separation between genotypes and treatment time points. Each point represents a biological replicate. (**C-G**) Gene Ontology (GO) enrichment analysis of DEGs in clusters 1–5, respectively. Top enriched GO terms for biological processes are shown for each cluster, with dot size indicating gene count and color scale indicating adjusted *p*-value. The arrows indicate the auxin-responsive GO terms.

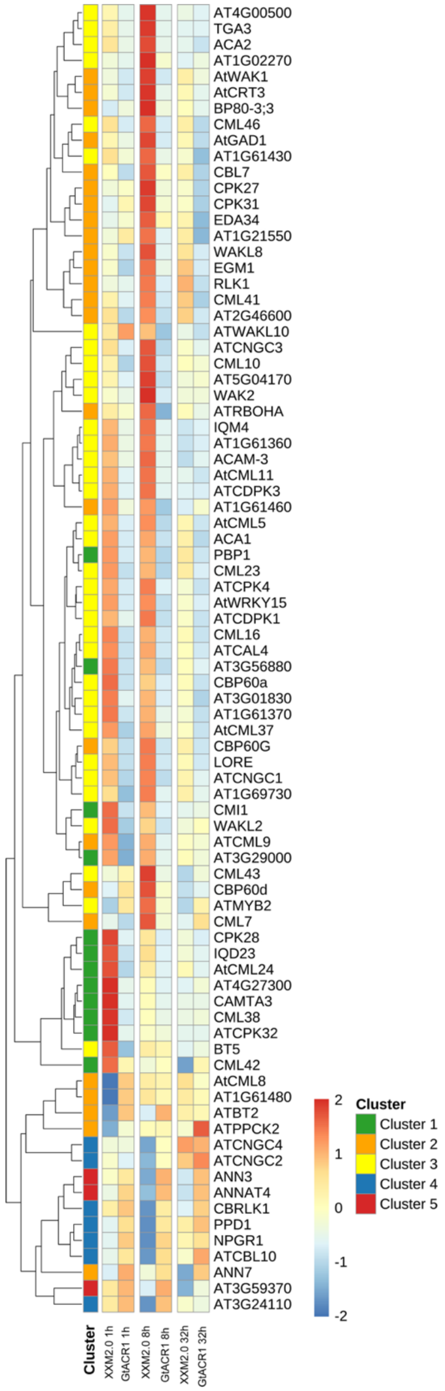


**Fig. S7. XXM2.0 activation triggers transcriptional responses of $Ca^{2+}$-responsive genes.**
Heatmap showing log2 fold changes of $Ca^{2+}$-associated genes in XXM2.0 and GtACR1 roots at 1 h, 8 h, and 32 h after blue light onset. Gene expression was clustered into five groups based on expression dynamics.

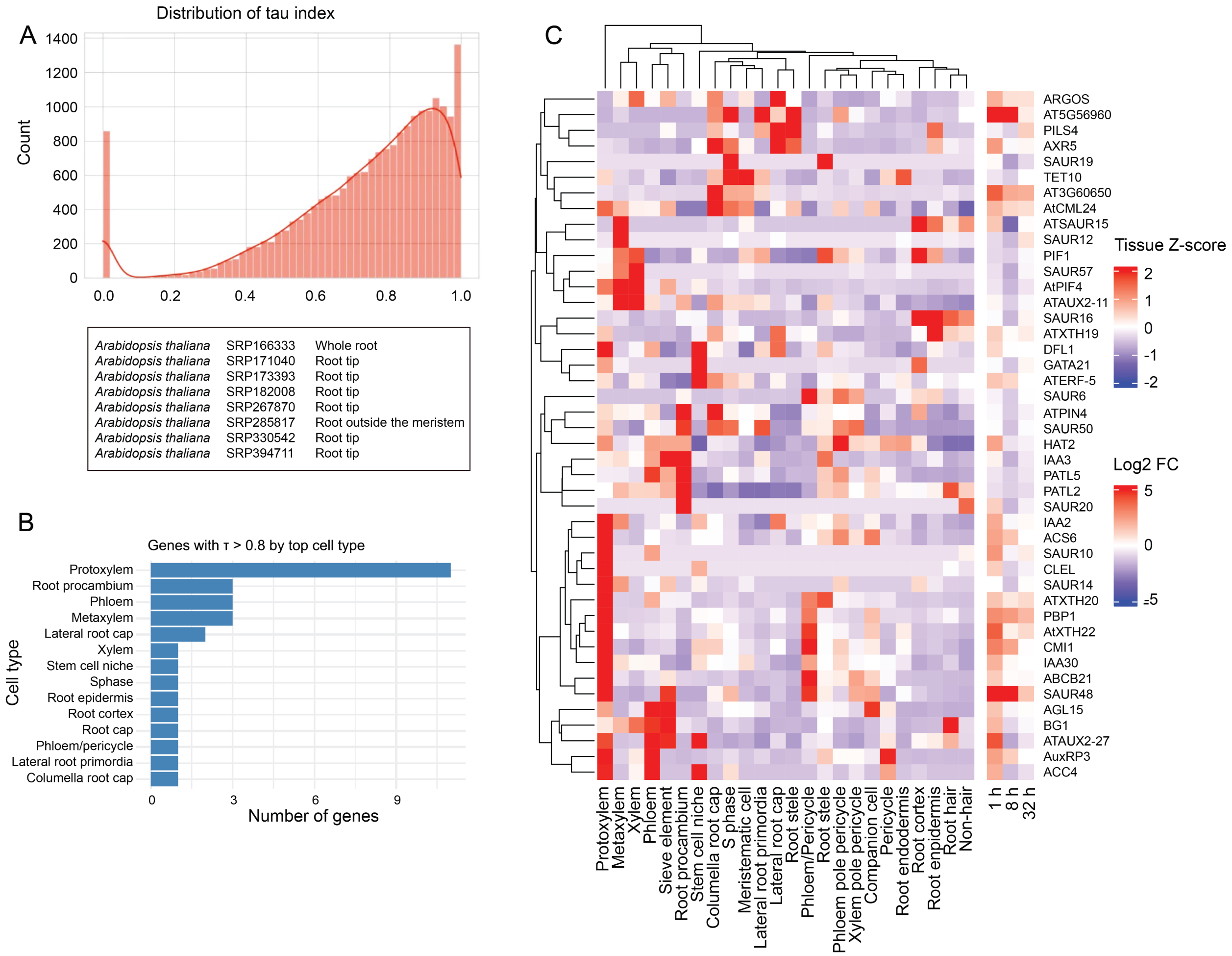


**Fig. S8. Identification of $Ca^{2+}$-sensitive tissues and cell types in Arabidopsis seedling roots.** (**A**) Distribution of the tau index for gene expression specificity across root cell types based on the Arabidopsis root atlas. Inset shows the top datasets used for root tissue profiling. (**B**) Bar plot showing the number of auxin-associated genes with high tissue specificity ($\tau > 0.8$) across different root cell types. The protoxylem, root procambium, and phloem were among the most enriched cell types. (**C**) Heatmap displaying the tissue-specific expression patterns of 45 auxin-associated DEGs from clusters 1, 4, and 5 identified in the XXM2.0 transcriptomic dataset (Fig. 4D). Expression data are mapped onto single-cell transcriptomes from the Arabidopsis root atlas. 36 of these genes exhibited strong enrichment (Z-score >1.5) in specific tissues or cell types. Color scale indicates normalized expression (Z-score) and log2 fold change (right margin, blue to red).

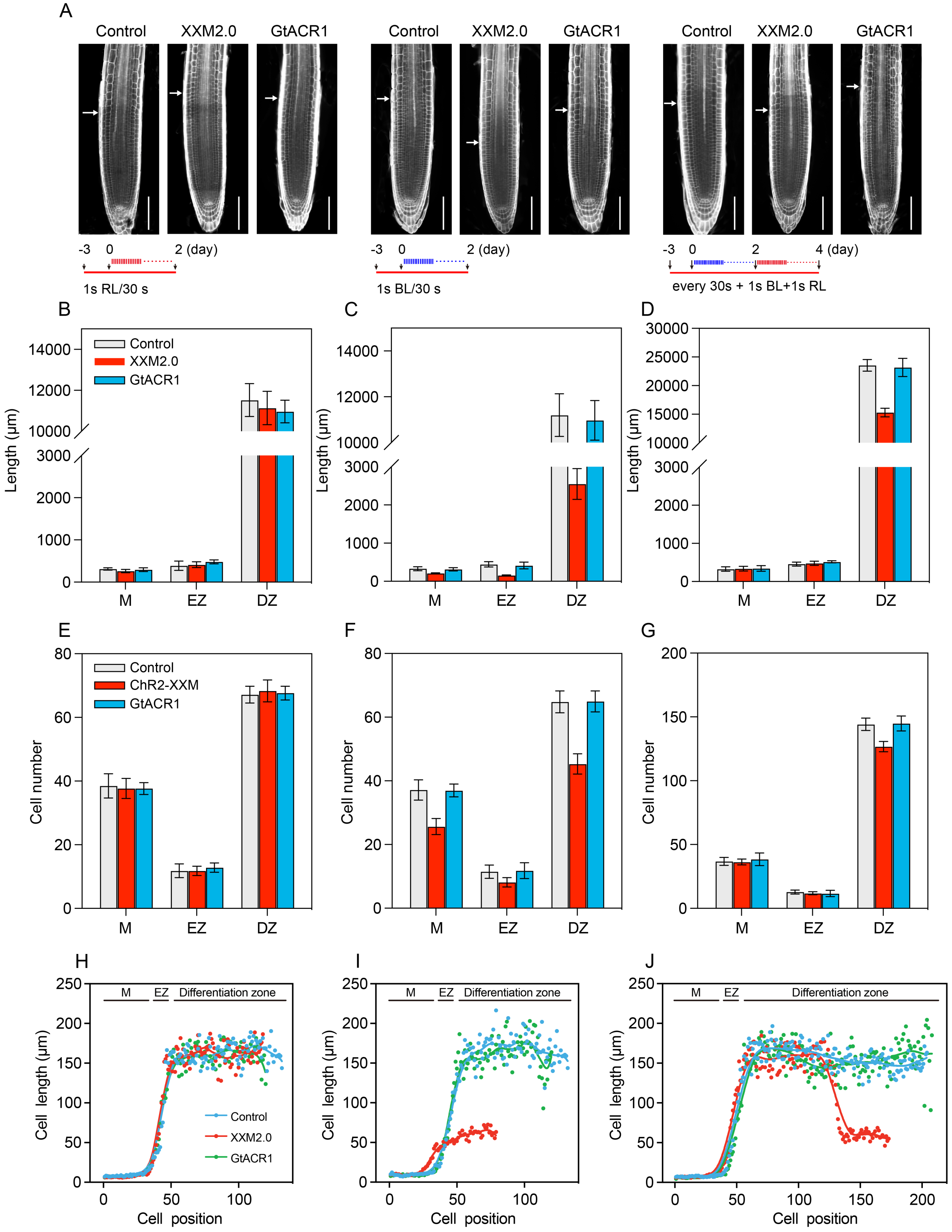
A
Control
XXM2.0
GtACR1
-3 0 2 (day)
1s RL/30 s
1s BL/30 s
-3 0 2 4 (day)
every 30s + 1s BL+1s RL
B
C
D
Control
XXM2.0
GtACR1
Length (μm)
M
EZ
DZ
E
F
G
ChR2-XXM
Cell number
H
I
J
M
EZ
Differentiation zone
Cell length (μm)
Cell position

1101

**Fig. S9. Spatial analysis of root zonation and cell number upon optogenetic stimulation.**
(**A**) Representative confocal images of root tips from control, XXM2.0, and GtACR1 seedlings subjected to different light treatments. Cell walls were stained with Calcofluor White to visualize cortical cell files. Roots were imaged under three conditions: RL or BL (1 s pulse every 30 s), or a switch from BL to RL after 2 days (recovery). The time axis is shown in days relative to light onset. Scale bar = 100 µm. (**B-D**) Quantification of cortex cell length in the meristem (M), elongation zone (EZ), and differentiation zone (DZ) after 2 days of either RL (C) or BL treatment (D), or a switch from BL to RL after 2 days (E). Error bars indicate standard error (n = 6). (**E-G**) Quantification of cortex cell numbers in the meristem, elongation zone, and differentiation zone after 2 days of either RL (E) or BL treatment (F), or a switch from BL to RL after 2 days (G). Error bars indicate standard error (n = 6). (**H-J**) Cell length in consecutive cortex cell files of Control, XXM2.0, and GtACR1 seedlings under pulsed light: 2 d RL (H), 2 d BL (I), and 2 d BL + 2 d RL (J). Measurements were taken from the quiescent center (position 1) to the light stimulation onset site, spanning the meristem (M), elongation zone (EZ), and differentiation zone. Individual dots represent mean cell length (n = 6), and lines show locally smoothed trends (LOWESS). Bars show mean ± SE, * $p < 0.05$, ** $p < 0.01$, *** $p < 0.001$.

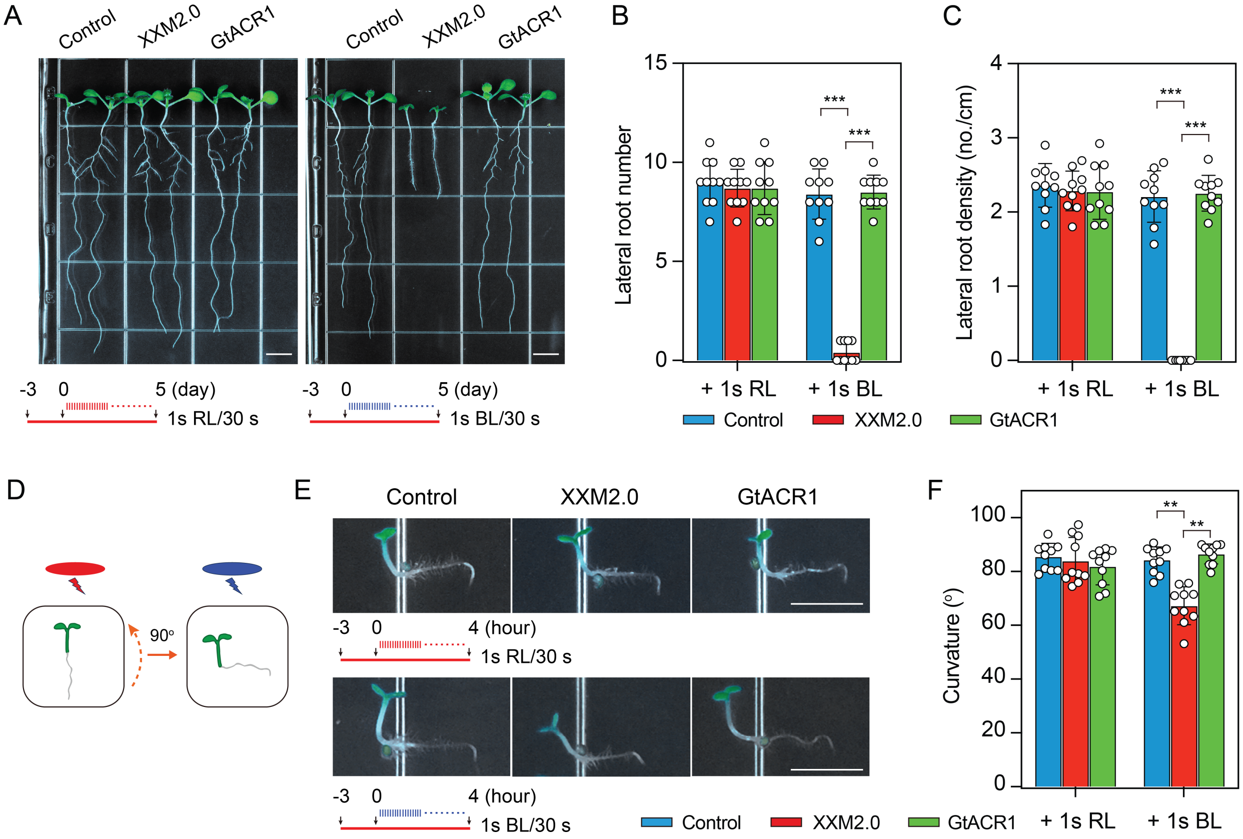


**Fig. S10. Optogenetically imposed $Ca^{2+}$ signals compromise lateral root formation and root gravitropism.**
(A) Representative lateral root phenotypes of 8-day-old transgenic Arabidopsis seedlings (Retinal only control, XXM2.0, and GtACR1). Seedlings were grown under continuous red LED illumination (650 nm, 100 µmol $m^{-2}$ $s^{-1}$) for 3 days, followed by 5 days of RL or BL pulse stimulation (1 s BL or RL, 30 s interval). Scale bars, 5 mm. (B–C) Quantification of lateral root number (B) and lateral root density (C) after 5 days of RL or BL pulse stimulation. Data represent mean ± SD (n = 10). *** $p < 0.001$. (D) Schematic diagram of the gravitropism assay. Seedlings were grown vertically on ½ MS medium under continuous red light for 3 days, after which plates were rotated 90° counterclockwise and maintained under RL or BL pulse stimulation for 4 h. (E) Representative gravitropic phenotypes of Control, XXM2.0, and GtACR1 seedlings following 90° reorientation. Scale bars, 3 mm. (F) Quantification of root curvature after 4 h of RL or BL pulse stimulation. Data represent mean ± SD (n = 10). ** $p < 0.01$.

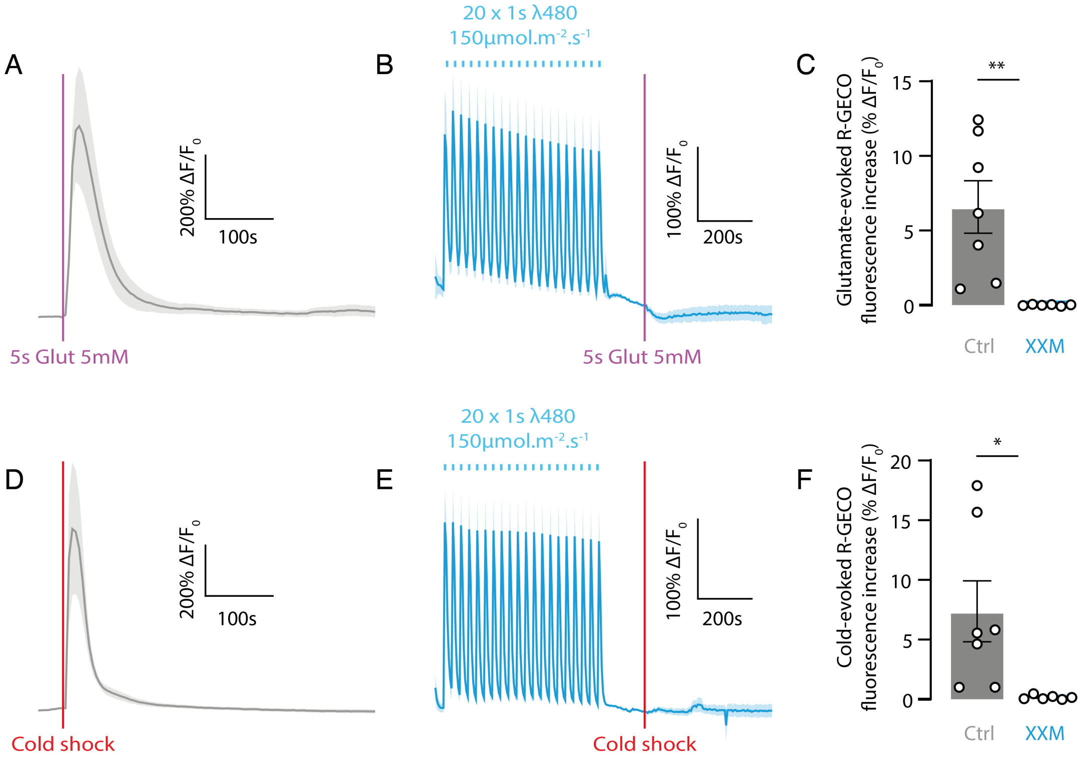


**Fig. S11 Optogenetically imposed $Ca^{2+}$ signals suppress glutamate- and cold-induced $Ca^{2+}$ responses.**
(A, D) Representative R-GECO1 fluorescence traces showing cytosolic $Ca^{2+}$ responses induced by local application of 5 mM glutamate (5 s) (A) or cold shock (D) in Arabidopsis root meristematic cells. (B, E) Representative R-GECO1 fluorescence traces showing glutamate- (B) or cold-induced (E) $Ca^{2+}$ responses following XXM2.0 activation by 20 × 1 s BL pulses ($\lambda = 480$ nm, 150 µmol $m^{-2}$ $s^{-1}$). (C, F) Quantification of glutamate-evoked (C) and cold-evoked (F) R-GECO1 fluorescence increases ($\Delta F/F_0$) in control and XXM2.0-stimulated roots. Data represent mean ± SD. Each dot indicates one biological replicate. * $p < 0.05$, ** $p < 0.01$.

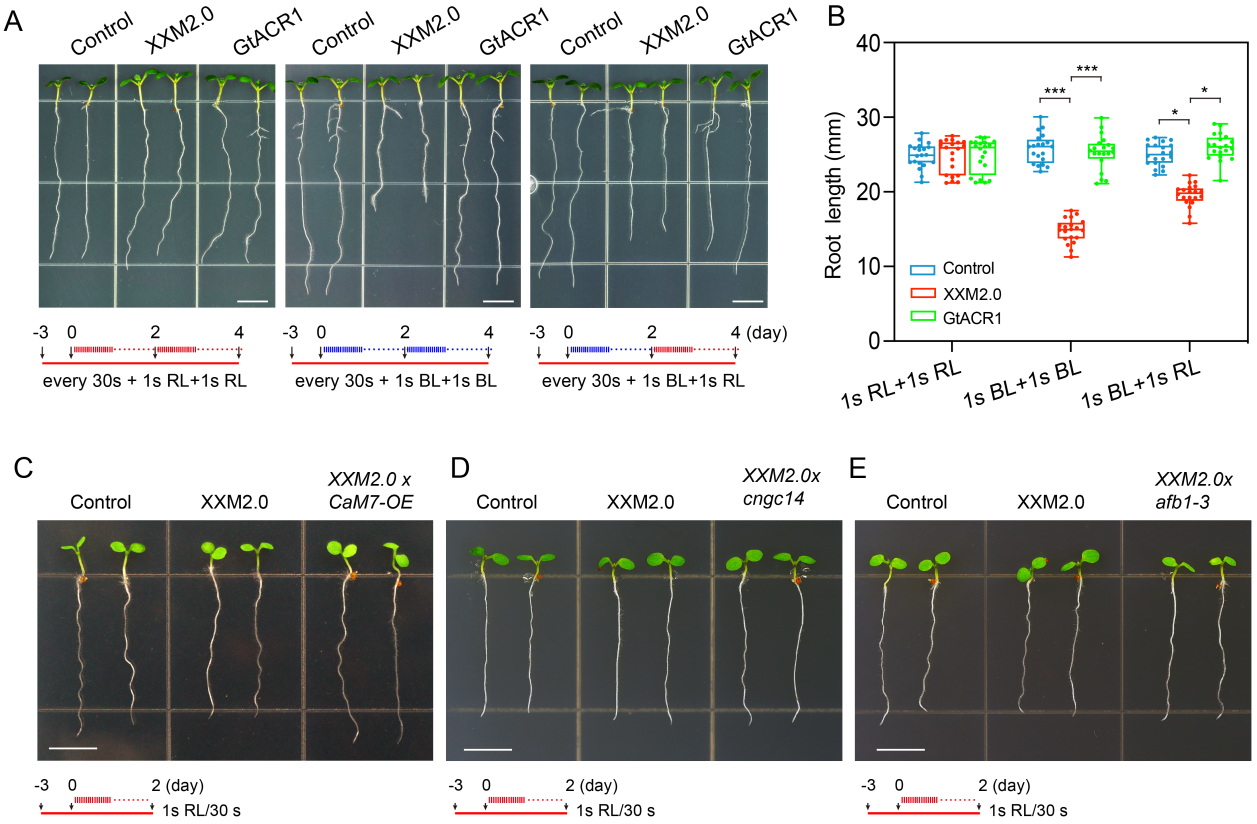


**Fig. S12 Reversible optogenetic control of root growth, and XXM2.0 phenotypes in CaM7-OE, *cngc14* and *afb1-3* backgrounds under RL**

(**A**) Root length phenotypes of 7-day-old Control, XXM2.0, and GtACR1 seedlings under three pulsed light treatments: 2 d RL + 2 d RL, 2 d BL + 2 d BL, and 2 d BL + 2 d RL. Scale bars = 5 mm. (**B**) Quantification of root length from (A). ($n$ = 20). *** $p < 0.001$, * $p < 0.05$. **(C-E)** Root length phenotypes of 5-day-old transgenic Arabidopsis lines Control, XXM2.0, and XXM2.0 × CaM7-OE (C), XXM2.0 × *cngc14* (D), or XXM2.0 × *afb1-3* (E). Seedlings were grown under continuous red LED illumination (650 nm, 100 µE) for 3 days, and followed by stimulation with red light (RL) pulses (1 s every 30 s) for an additional 2 days. Scale bars = 5 mm.

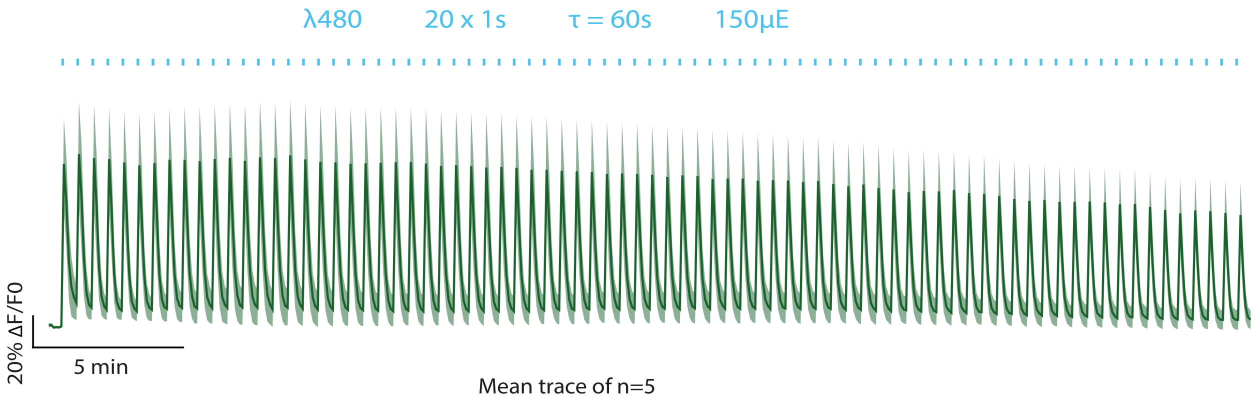


**Fig. S13. Cytosolic $Ca^{2+}$ dynamics during prolonged XXM2.0 photostimulation.** Representative mean R-GECO1 fluorescence trace showing cytosolic $Ca^{2+}$ signals during 30 min of repetitive XXM2.0 activation using 20 × 1 s blue-light pulses (λ = 480 nm, τ = 60 s, 150 µmol $m^{-2}$ $s^{-1}$). Shaded area indicates ± SD. Mean trace of n = 5 biological replicates.

### Table S1. List of primers used in this study

| Primer name | Sequence (5'-->3') |
|---|---|
| ChR2-XXM2.0 USER fwd | GGCTTAAUCGCGAGCTGCTATTTGTAAC |
| ChR2-XXM2.0 USER oS rev | GGTTTAAUCCCTCGACTACCGCGCCAG |
| R-GECO1 USER fwd | GGTTTAAUATGGTCGACTCTTCACGTCGTA |
| R-GECO1 USER rev | GGTTTAAUCTACTTCGCTGTCATCATTTGTA |
| R-GECO1 linker overlap fwd | AAACCTCAGUGGTGGAATGGTCGACTCTTCACGTCGT |
| R-GECO1 linker overlap rev | ACTGAGGTTUAATCCCTTCGCTGTCATCATTTGTACA |
| CaM7 USER fwd | GGCTTAAUATGGCGGATCAGCTAACC |
| CaM7 USER oS rev | GGTTTAAUCCCTTTGCCATCATGACTTTG |

**Table S2. List of datasets used for tissue-specific expression analysis in the root**

| Dataset | Bioproject | Species | Tissue | Libraries | Age | Experiments | Cells | PMID |
|---|---|---|---|---|---|---|---|---|
| SRP166333 | PRJNA497883 | Arabidopsis thaliana | Whole root | 10x Genomics | 7 days old seedling | 3 | 16949 | 30923229 |
| SRP171040 | PRJNA507252 | Arabidopsis thaliana | Root tip | 10x Genomics | 5 days after germination | 5 | 33956 | 30718350 |
| SRP182008 | PRJNA517021 | Arabidopsis thaliana | Root tip | 10x Genomics | 10 days old seedling | 1 | 13514 | 31004836 |
| SRP285817 | PRJNA666436 | Arabidopsis thaliana | Root outside the meristem | 10x Genomics | 4-5 days after germination | 7 | 17553 | 33822225 |
| SRP330542 | PRJNA750934 | Arabidopsis thaliana | Root tip | 10x Genomics | 6 days old seedling | 2 | 22606 | 34358681 |
| SRP173393 | PRJNA509920 | Arabidopsis thaliana | Root tip | 10x Genomics | 6 days old seedling | 3 | 13252 | 30913408 |
